# Chemoproteomics identifies a pyrimido[4,5-*d*]pyrimidine analogue as a tubulin-tyrosine ligase binder

**DOI:** 10.64898/2026.08.06.743276

**Authors:** Rubaba R. Abanti, Eleftheria A. Georgiou, Dmytro Makarov, Severin Lechner, Afroditi Tsigara, Bernhard Kuster, Guillaume Médard, Pavel Kielkowski, Leentje Persoons, Steven De Jonghe, Ioannis K. Kostakis

## Abstract

Small-molecule drug discovery relies on identifying compounds that modulate specific protein targets, a process often hindered by cellular complexity. Through phenotypic screening of a kinase-focused diazaquinazoline library, we serendipitously identified **CEM198** as the first high-affinity ligand of tubulin-tyrosine ligase (TTL). Functional assays combining live-cell TTL inhibition, microtubule polymerization, cell cycle analysis, and proteomics revealed that **CEM198** acts through a dual mechanism: directly binding to TTL and altering α/β-tubulin conformation. This interaction restricts α-tubulin tyrosination and disrupts tubulin polymerization, leading to microtubule destabilization. The differential effects observed between SH-SY5Y and HEK293T cells indicate that effective TTL inhibition depends on both direct binding and structural modulation of the tubulin heterodimer. These findings introduce **CEM198** as a chemical probe for investigating the tubulin tyrosination–detyrosination and demonstrate the potential of chemoproteomics to uncover novel modulators of microtubule dynamics.

## Introduction

Small molecule drug discovery traditionally hinges on identifying compounds that modulate the activity of specific protein targets.^[1,2]^ This process often starts with phenotypic screening, followed by strategies to delineate the mode-of-action that links a compound to its target and the resulting phenotype. However, the inherent complexity of cellular systems and protein networks renders this process highly challenging. Chemoproteomic methods have revolutionized the identification of protein targets for bioactive small molecules.^[3–5]^ Enabled by advances in quantitative mass spectrometry, chemoproteomics now facilitates high-throughput screening of compound libraries to identify protein targets.^[6–8]^ It has been successfully applied to determine protein binders of diverse natural products and therapeutic agents, including sorafenib^[9]^, eupalmerin^[10]^ and artemisinin.^[11]^ The core strategy involves the pull-down of interacting proteins using the bait compound, followed by the protein identification utilizing liquid chromatography–tandem mass spectrometry (LC-MS/MS).^[4,12–14]^ Furthermore, the affinity of the small molecule-protein complex can be probed by competition assays with the parent compound, in which increasing concentrations of the free (non-immobilized) parent compound displace target proteins from the bait immobilized on beads. This approach allows a high degree of parallelization and automation, enabling screens of large compound libraries, such as those composed of kinase inhibitors.^[7]^ In this article, we report on the phenotypic antitumoral screening of (diaza)-quinazoline derivatives, which led to the serendipitous discovery of a small molecule modulator of microtubules stability.

Microtubules are crucial components of the cytoskeleton, involved in cell division, polarity, migration and intracellular vesicle transport.^[15]^ Dysregulation of microtubule dynamic is a hallmark of various pathological conditions, including cancer and neurodegenerative diseases, making them attractive targets for therapeutic intervention.^[16–19]^ The stability of microtubules and their interactions with microtubule associated proteins (MAPs) are modulated by a series of post-translational modifications (PTMs), including acetylation, detyrosination/tyrosination, (poly)glutamylation, and (poly)glycylation, collectively referred to as the ‘tubulin code’.^[15,20]^ Although the enzymes catalyzing the introduction and removal of these PTMs have been identified, small molecule modulators capable of selectively targeting them remain largely unavailable. The methods to detect tubulin modifications traditionally relied on the use of modification-specific antibodies or the application of isotope-labeled substrates.^[19,21–23]^Recently, we introduced a chemoproteomic workflow to monitor tubulin tyrosination in living cells using a metabolically incorporated tyrosine propargyl probe (Tyr-*O*-Alk), which is enzymatically added to the C-terminus of α-tubulin by tubulin-tyrosine ligase (TTL).^[24]^ TTL exhibits low substrate specificity and is therefore capable of incorporating several unnatural tyrosine analogues, including Tyr-*O*-Alk.^[24]^TTL specifically modifies soluble tubulin dimers, which are then assembled into new microtubules that can subsequently be stabilized by detyrosination.^[25]^ TTL features an ATP binding pocket, but has not been reported to be inhibited by any kinase inhibitor. Interestingly, the first small molecule inhibitors of TTL activity were identified among a group of sesterterpenes isolated from *Salvia dominica*.^[26,27]^ Data from the Mouse Phenotyping Consortium, indicate that TTL knockout (KO) in mice leads to edema, reduced exploratory behavior and pre-weaning lethality, highlighting its physiological importance.^[26,28]^

Here, we report a phenotypic anticancer screening of a kinase-focused library containing (diaza)-quinazoline derivatives, which unexpectedly led to the identification of the first in class high-affinity TTL ligand – **CEM198** (**Figure 1A**). Functional characterization of **CEM198** using live-cell TTL inhibition assays, microtubule polymerization analysis, cell cycle profiling and proteomics revealed dual activity of the compound by restricting α-tubulin tyrosination and tubulin polymerization.

**Figure 1.**
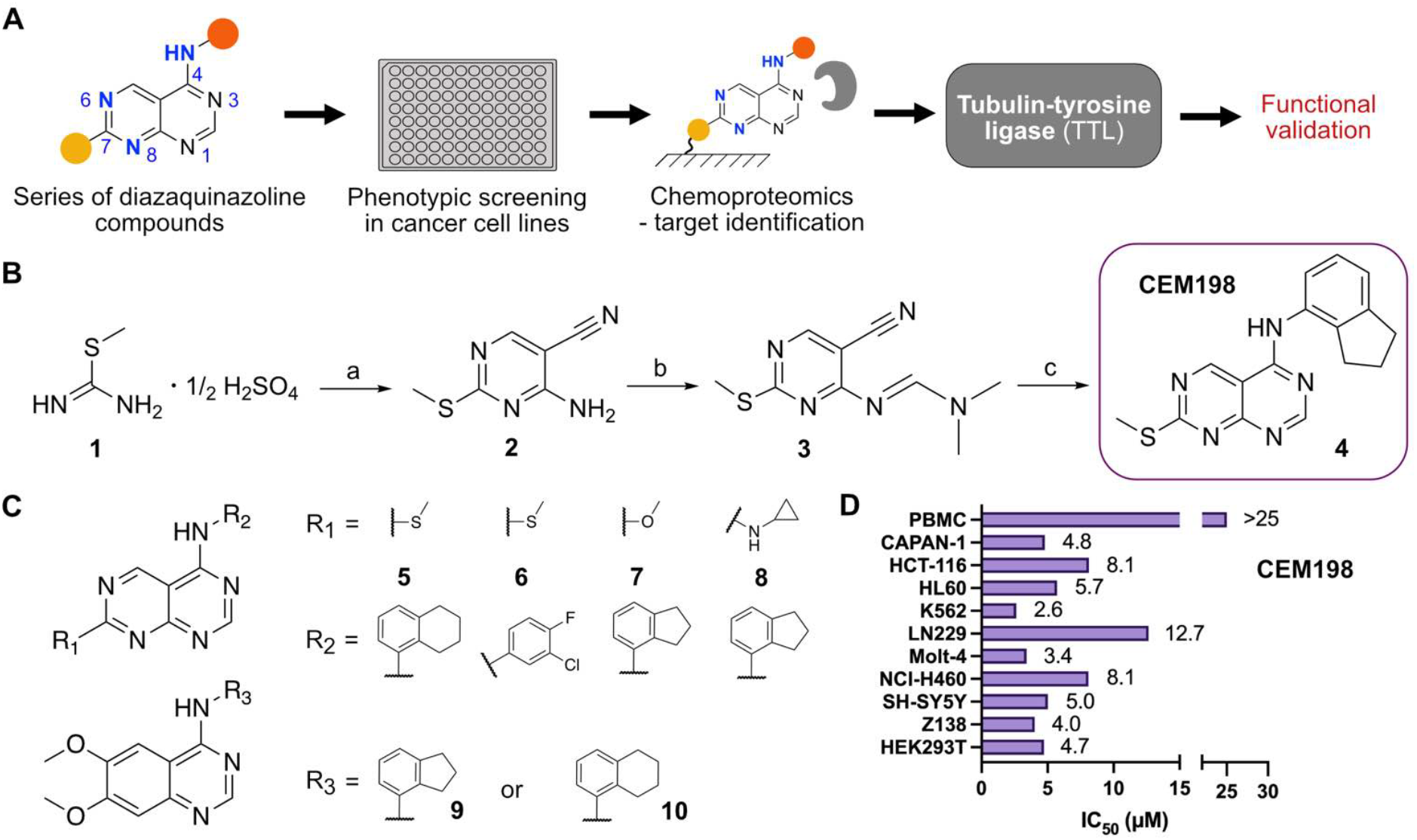
Overview of **CEM198** characterization as TTL binder. **A.** The workflow leading to identification of TTL specific binder **CEM198. B**. Synthesis and structure of **CEM198**; (a) 2-(ethoxymethylene)malononitrile, triethylamine, ethanol abs., rt; (b) DMF-DMA, toluene, reflux; (c) appropriate aniline, CH_3_CO_2_H, reflux. **C**. Structures of synthetized and tested compounds. **D**. Bar chart of IC_50_ (μM) values across ten cancer cell lines and normal peripheral blood mononuclear cells (PBMCs) after 72 hours of treatment.

## Results

### Identification of CEM198 as a potent and selective antiproliferative agent

Building on our previous screening efforts aimed at discovering novel anticancer agents via potential kinase inhibition, we designed and synthesized a series of pyrimido[4,5-*d*]pyrimidine fanalogs (**Figure 1A**). These compounds share key features with 4-anilinoquinazolines, a well-known class of EGFR inhibitors.^[29]^ In particular, the new analogs possess two nitrogen atoms at positions 1 and 3, which are critical for activity due to their ability to form essential hydrogen bonds within the ATP-binding pocket. A notable structural difference from 4-anilinoquinazolines is the presence of two additional nitrogen atoms at positions 6 and 8 (**Figure 1A**). The nitrogen at position 6 occupies a site where other known EGFR inhibitors typically bear small, electron-donating substituents. Moreover, the new derivatives feature an amino side chain at position 7, replacing the alkoxy group commonly found in related scaffolds.^[29]^ Based on the preliminary results and the promising nature of this scaffold, we synthesized a new set of analogs—along with several anilinoquinazoline derivatives for comparison—to further explore and optimize their biological activity.

The synthesis of the pyrimido[4,5-*d*]pyrimidines is outlined in **Figure 1B** and Scheme S1, whilst the synthesis of the quinazoline analogs is depicted in Scheme S2. Initially, the sulfate salt of 2-methyl-2-thiopseudourea (**1**) was prepared by treatment of thiourea with dimethyl sulfate at 80°C. Subsequently, the reaction between 2-(ethoxymethylene)malononitrile and derivative **1**, in the presence of triethylamine, afforded the desired pyrimidine **2**, which, upon reaction with *N,N*-dimethylformamide dimethyl acetal (DMF-DMA) yielded intermediate **3**. Finally, the desired pyrimido[4,5-*d*]pyrimidines **4, 5** and **6** were obtained upon treatment with the appropriate aniline (**Figure 1B, C**). The methyl thioethers **4**-**6** were complemented by corresponding methoxy (**7**) and cyclopropylamine (**8**) derivatives, prepared via an analogous sequence involving oxidation of the methylthio group (**Figure 1C**, Scheme S1). In parallel, two quinazoline analogs **9** and **10** were synthesized for comparison (**Figure 1C**, Scheme S2).

The compounds were evaluated for anti-proliferative activity across a panel of nine cancer cell lines, including pancreatic adenocarcinoma (Capan-1), colorectal carcinoma (HCT-116), acute myeloid leukemia (HL-60), chronic myeloid leukemia (K-562), glioblastoma (LN-229), acute lymphoblastic leukemia (Molt-4), lung carcinoma (NCI-H460), neuroblastoma (SH-SY5Y), non-Hodgkin lymphoma (Z-138), and the human embryonic kidney cell line HEK293T, which was included for mechanistic deconvolution rather than for cancer profiling. Cells were treated for 24 and 72 hours (**Figure 1D** and Table S1). While most derivatives showed moderate anti-proliferative activity, compound **4**, hereafter referred as **CEM198**, stood out as highly potent, with IC50 values ranging from 2.6 to 8 μM depending on the cell line. K562 cells were particularly sensitive, with an IC50 of 2.6 μM. To assess selectivity, **CEM198** was also tested against normal, non-cancerous peripheral blood mononuclear cells (PBMCs) and showed no cytotoxicity up to 25 μM (**Figure 1D**, Table S1), indicating that its anti-proliferative effect is largely selective for cancer cells.

### Target deconvolution of CEM198

Although **CEM198** emerged as a potent and selective antitumoral agent in phenotypic assays, the molecular target responsible for its cytotoxicity remained unknown. Therefore, to elucidate the molecular target(s) underlying this phenotype, a kinobeads-based chemoproteomic profiling was performed allowing the determination of the binding affinity of **CEM198** to approximately 200 kinases expressed in K562 cell lysates (**Figure 2A**).^[30]^ Surprisingly, **CEM198** did not exhibit measurable affinity to any of the kinases covered by the kinobeads platform. Instead, **CEM198** showed dose-dependent binding to dCTP pyrophosphatase 1 (DCTPP1, Kdapp=1.9 µM), a protein previously identified as an off-target for several kinase inhibitors, including GSK-690693 and Milciclib.^[31]^ However, since DCTPP1 is not classified as an essential gene in the DepMap dataset^[32]^ encompassing over 900 cancer cell lines, we hypothesized that additional non-kinase targets, outside the coverage of the kinobeads assay, might be responsible for the cytotoxic effects of **CEM198**.

**Figure 2.**
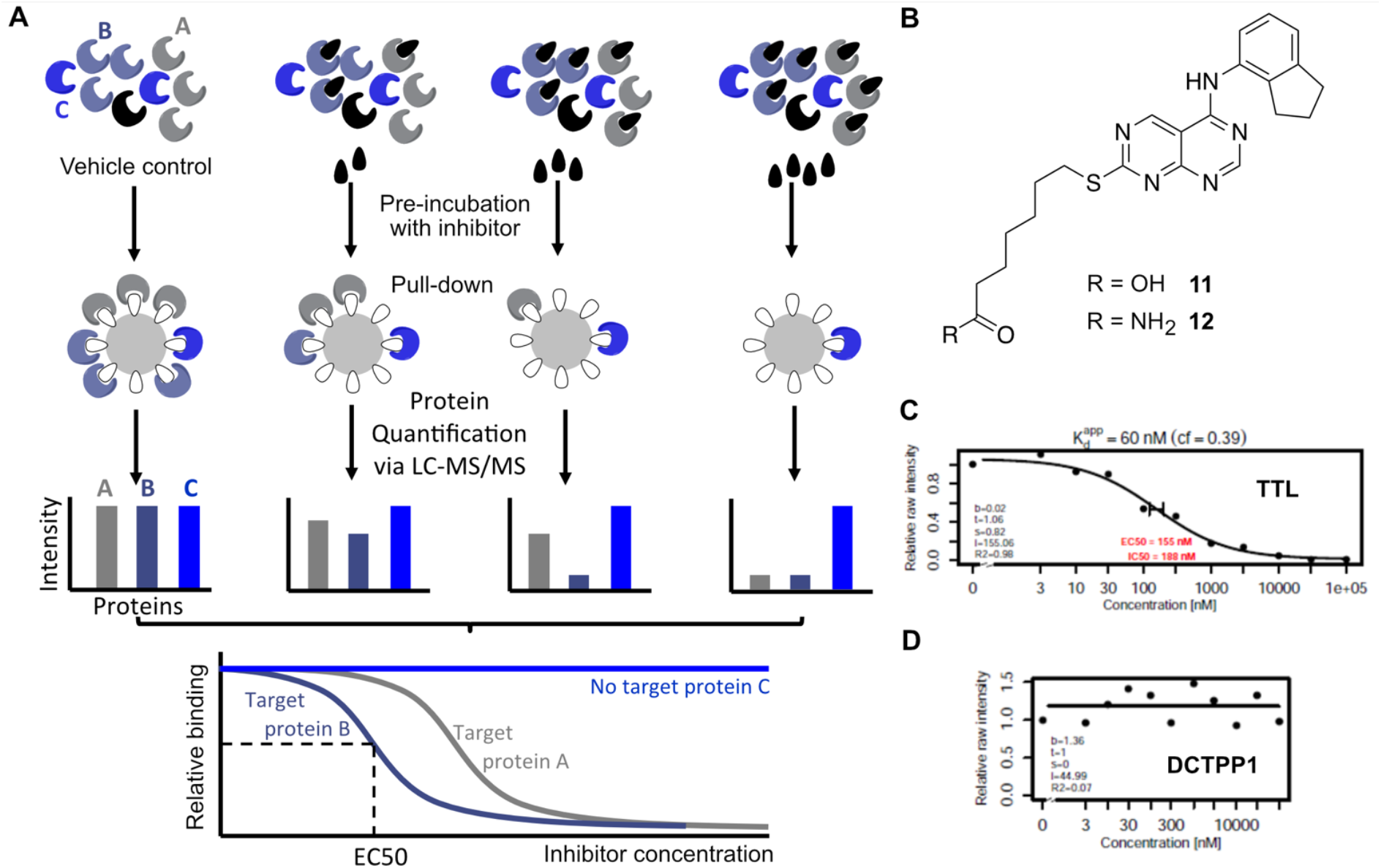
Chemoproteomics screening identifies **CEM198**-TTL interaction. **A.** Scheme of chemoproteomics target deconvolution. **B**. Structures of compounds **11** and **12** used for chemoproteomics pull-down of **CEM198** protein targets. **C. CEM198** concentration dependent TTL pull-down from K562 cell lysate. **D**. DCTPP1-**CEM198** affinity – control.

To identify such targets, we designed a chemoproteomics competition pull-down assay tailored to **CEM198** (**Figure 2A**), enabling detection of specific protein interactions in a cellular context. For this purpose, we synthesized the pyrimido[4,5-*d*]pyrimidine derivative **11**, incorporating a linker at the sulfur atom and equipped with a terminal carboxylic acid to enable immobilization on Sepharose beads (**Figure 2B** and Scheme S3). Because the introduction of a linker could potentially interfere with the biological activity of the scaffold, and since the negatively charged carboxylic acid in **11** might impair cellular permeability, we also prepared the corresponding carboxamide analog **12** (Scheme S3). Both compounds **11** and **12** were evaluated for antitumoral activity to ensure that the structural modifications did not compromise their biological function. Both derivatives retained anticancer activity against the same panel of cancer cell lines, including K562 cells used for chemoproteomic experiments, suggesting that they remained capable of engaging the relevant cellular target (Table S1). In the chemoproteomic competition assay, lysates of K562 cells pretreated with increasing concentrations of **CEM198** were incubated in the presence of immobilized **11**. Proteins bound by **CEM198** were prevented from being pulled down by the immobilized **11**, resulting in a dose-dependent reduction in the signal in the mass spectrometry readout. Only one protein, TTL, showed dose-dependent competition, yielding an EC_50_ of approximately 150 nM (**Figure 2C**). Notably, DCTPP1, previously detected in kinobeads profiling, was not enriched in this assay, supporting TTL as the primary and specific target of CEM198 (**Figure 2D**). Given that no small-molecule ligands for TTL have been reported and that TTL features an ATP-binding pocket, this finding is particularly compelling. Although it remains to be determined whether TTL inhibition alone accounts its cytotoxicity, these results clearly establish TTL as a high-affinity target of CEM198 in K562 cells.

### Cellular target engagement assay

To confirm cellular engagement of TTL by **CEM198**, we employed a previously established tyrosination assay. TTL catalyzes the incorporation of a C-terminal tyrosine residue onto α-tubulin (TUBA).^[24]^ The reverse reaction, i.e. detyrosination, is performed by tubulin-Tyr carboxypeptidase 1 and 2 (VASH1 and 2) in complex with small vasohibin-binding protein (SVBP) or microtubule-associated tyrosine carboxypeptidase 1 (MATCAP, **Figure 3A**).^[33,34]^ The tyrosination assay is based on the TTL-catalyzed metabolic incorporation of the tyrosine analog probe Tyr-*O*-Alk into α-tubulin in the neuronal-like SH-SY5Y cells, which exhibit robust amounts of microtubule modifications (**Figure 3B**).^[24]^ Upon incorporation into TUBA, the terminal alkyne of the Tyr-*O*-Alk is coupled with rhodamine-azide using a Cu(I)-catalyzed azide-alkyne cycloaddition (CuAAC) ‘click chemistry’.^[12,35,36]^ The protein mixture is then separated by sodium dodecyl-sulfate polyacrylamide gel (SDS-PAGE) and the resulting fluorescent signal is quantified to assess TUBA labelling. Initial experiments revealed that prolonged exposure (72 hours) to **CEM198** at concentrations ranging from 25 nM to 30 μM induced significant cytotoxicity and cell detachment in SH-SY5Y cells (IC_50_ 5 μM, Table S1).

**Figure 3.**
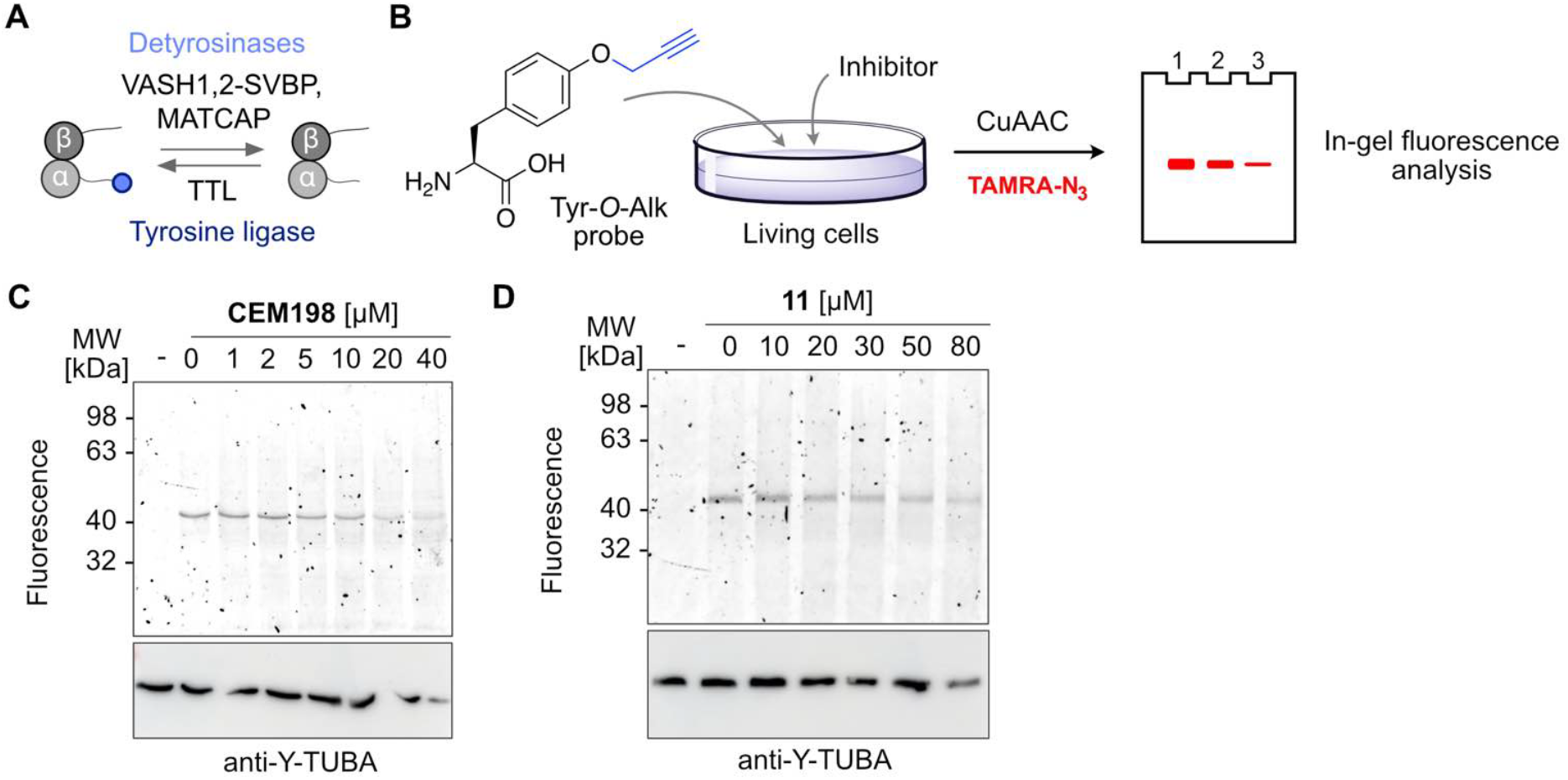
Tubulin tyrosination-detyrosination cycle inhibition in SH-SY5Y cells. **A.** Scheme of the tubulin tyrosination-detyrosination cycle with involved enzymes. **B**. Scheme of the TTL activity assay using Tyr-*O*-Alk probe. **C**. Concentration-dependent TTL inhibitory activity of **CEM198** in SH-SY5Y cells. **D**. Concentration-dependent TTL inhibitory activity of bait compound **11** in SH-SY5Y cells. For **C** and **D** the cells were treated with Tyr-*O*-Alk probe and inhibitor together for 24 h. Western blot of the corresponding SDS-PAGE after fluorescence scan with antibody recognizing tyrosinated α-tubulin.

Reducing treatment duration from 72 to 24 hours partially mitigated these effects (Table S1). The concentration-dependent treatment with **CEM198** of SH-SY5Y cells resulted in an inhibition of TLL-mediated incorporation of the Tyr-*O*-Alk probe, with near complete inhibition at a concentration of 20 µM (**Figure 3C**). In contrast, immunostaining for tyrosinated α-tubulin showed only modest reduction (**Figure 3C**). This is likely due to the limited turnover of detyrosinated α-tubulin within the 24-hour window and the large difference in absolute levels of tyrosinated versus detyrosinated α-tubulins. Similarly, the bait compound **11** (**Figure 3D**), utilized in the chemoproteomic profiling, also inhibited TTL activity in a concentration dependent manner (**Figure 3D**). To further explore structure–activity relationships (SAR), we evaluated a series of the **CEM198** analogu, including **5**-**10** (Figure S1-S6). They exhibited minimal or no TTL inhibition, highlighting the importance of the aminoindane moiety of the **CEM198** scaffold for target engagement. Collectively, these results support the conclusion that both **CEM198** and **11** inhibit TTL catalytic activity in SH-SY5Y cells.

To further validate TTL as a functional target of **CEM198**, TTL was transiently overexpressed in HEK293T cells, and its enzymatic activity was monitored using the Tyr-*O*-Alk incorporation assay (**Figure 4A**). TTL overexpression led to a mild increase in tyrosinated TUBA and a corresponding reduction in detyrosinated α-tubulin, as confirmed by Western blot analysis with modification-specific antibodies, consistent with enhanced TTL activity (**Figure 4A**). MS-based proteomics confirmed overexpression of the TTL-GFP fusion protein and revealed only minimal changes in the global proteome (Figure S7). Additionally, Tyr-O-Alk incorporation increased approximately two-fold compared to non-transfected cells, further supporting functional activity of the overexpressed enzyme (**Figure 4A**). Interestingly, treatment of TTL-overexpressing HEK293T cells with **CEM198** did not reduce Tyr-*O*-Alk incorporation, suggesting limited inhibition of TTL catalytic activity under these conditions (**Figure 4B**). However, a pronounced increase in detyrosinated α-tubulin was observed following treatment with 160 μM of **CEM198** (**Figure 4B**). This effect was also evident in non-transfected HEK293T cells, where **CEM198** treatment led to substantial accumulation of detyrosinated α-tubulin and cell detachment (**Figure 4C**). In contrast, other analogs, including **9**-**11**, did not elicit similar effects in HEK293T cells (Figure S8-S10). Notably, **12**, the primary amide analogue of the carboxylic acid **11**, led to a comparable increase in detyrosinated α-tubulin levels in both HEK293T cells with TTL overexpression (**Figure 4D**) and in detached non-transfected cells (**Figure 4E**). Although Tyr-O-Alk incorporation was not markedly affected by these treatments, likely due to the relatively low turnover of microtubule modifications in HEK293T cells, the observed increase in detyrosinated α-tubulin upon exposure to **CEM198** and **12** suggests a reduction in TTL activity. These findings suggest that while **CEM198** and its analogs may not uniformly inhibit TTL catalytic activity across cell types, they nonetheless perturb the tubulin tyrosination–detyrosination cycle.

**Figure 4.**
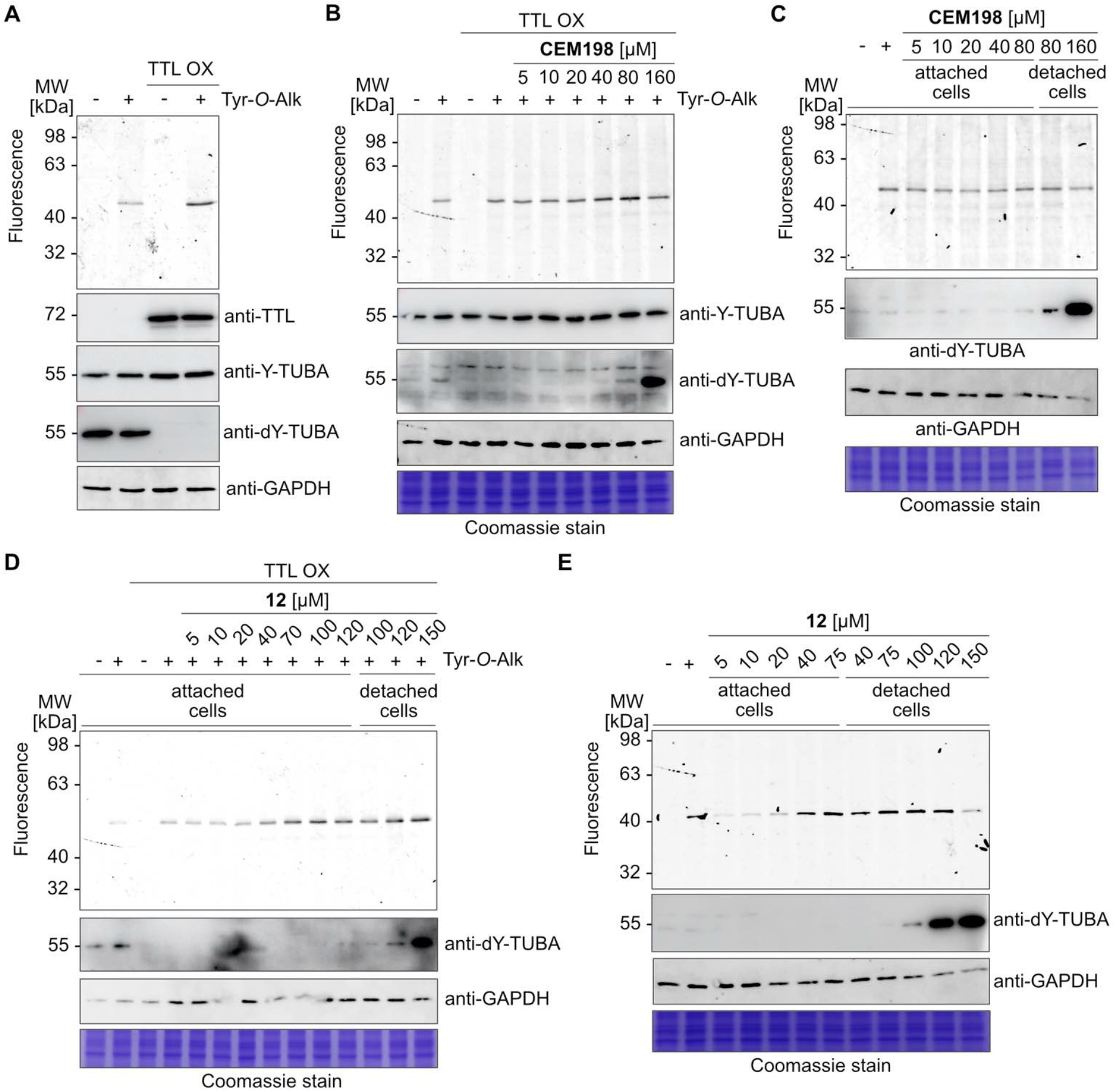
Inhibition of TTL in TTL overexpressing **HEK293T** cells. **A.** In HEK293T cells w/ and w/o TTL OX Tyr-*O*-Alk probe incorporation and changes in levels of tyrosinated/detyrosinated α-tubulin. **B**. Concentration dependent **CEM198** activity TTL OX HEK293T cells. **C**. Concentration dependent **CEM198** activity in HEK293T cells without TTL OX. **D**. Concentration dependent **12** activity in HEK293T cells with TTL OX. **E**. Concentration dependent **12** activity in HEK293T cells without TTL OX.

Treatment of both SH-SY5Y and HEK293T cells with **CEM198** resulted in pronounced cell detachment, particularly upon prolonged exposure (72 hours) or at higher concentrations (> 80 μM). This phenotype is consistent with microtubule disassembly. However, this effect is unlikely to be solely attributable to TTL inhibition, as increased levels of detyrosinated α-tubulin are generally associated with microtubule stabilization rather than destabilization. These observations suggest that **CEM198** may exert dual activity—both inhibiting TTL and interfering with microtubule assembly, potentially by disrupting α/β-tubulin heterodimer formation or inhibition of their polymerization into microtubules. To investigate this hypothesis, we assessed the impact of **CEM198** and **12** on microtubule polymerization *in vitro* using a fluorescence-based assay that monitors the real-time assembly of purified tubulin into microtubules, as indicated by an increase in fluorescence intensity (**Figure 5A**). Vinblastine and colchicine were included as positive controls, as both are well-characterized microtubule-destabilizing agents that inhibit polymerization by binding tubulin and preventing microtubule elongation. As expected, these controls produced a pronounced decrease in polymerization kinetics, validating the assay performance (Figure S11). Since both **CEM198** and **12** exhibit tubulin polymerization inhibitory activity, with **12** displaying markedly greater potency, we next compared their cellular effects to those of colchicine and vinblastine.^[37,38]^ Colchicine is a well-characterized microtubule destabilizer that binds to β-tubulin at the α/β interface, leading to inhibition of microtubule polymerization (Figure S12).^[39]^ Colchicine did not affect TTL-mediated tyrosination nor did induce cell detachment in HEK293T cells (Figure S11). In contrast, vinblastine, which binds to the α/β-tubulin heterodimer and induces a conformational change disfavoring microtubule assembly, caused both cell detachment and a significant increase in detyrosinated α-tubulin levels (**Figure 5B**).

**Figure 5.**
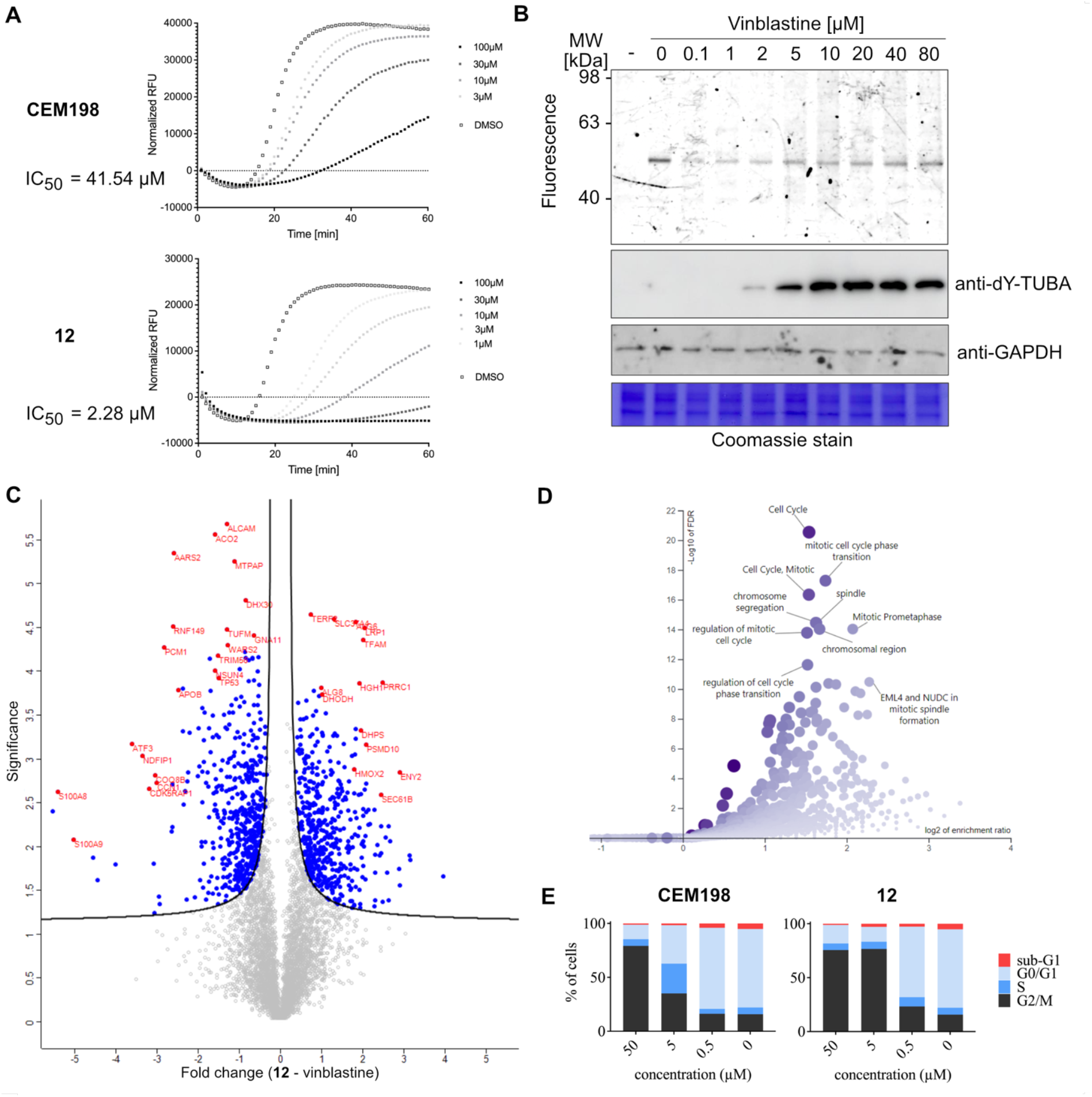
Evaluation of **CEM198** impact on microtubule polymerization and cell cycle. **A.** The effect of **CEM198** and **12** on microtubule polymerization *in vitro* with corresponding IC_50_ values. **B**. Probing the vinblastine activity in TTL tyrosination assay, complemented by Western blot using anti-detyrosinated α-tubulin to determine changes in tyrosination. **C**. Differences in whole proteome upon treatment of HEK293T cells with **12** or vinblastine; Significance is -log_10_(*p*-value), fold change is defined by log_2_(**12** – vinblastine). **D**. GO terms analysis of downregulated proteins in HEK293T cells treated with **12** in comparison to untreated control. **E**. Concentration dependent cell cycle arrest by **CEM198** or **12** in SH-SY5Y cells after 24 hours of treatment.

To further delineate the cellular effects of **12**, we performed comparative proteomic profiling of HEK293T cells treated with **12** and vinblastine (**Figure 5C**). The resulting data revealed distinct proteomic signatures (Figure S13). Gene ontology (GO) analysis of downregulated proteins in HEK293T cells treated with **12** points towards strong disruption of cell cycle progression and mitotic arrest (**Figure 5D**). These findings were corroborated by cell cycle analysis, which showed predominant arrest in the G2/M phase after treatment with **12**, in agreement with its action as a tubulin polymerization inhibitor (**Figure 5E**). Similarly, **CEM198** arrested cells in G2/M phase (**Figure 5E**), although at slightly higher concentrations, reflecting its weaker inhibition of tubulin polymerization.

Collectively, these findings suggest that both vinblastine and **CEM198** interfere with TTL mediated α-tubulin tyrosination, likely by inducing conformational changes in the α/β-tubulin heterodimer prohibiting efficient interaction with the TTL. Importantly, while **CEM198** binds TTL with high affinity, the differential effects observed between SH-SY5Y and HEK293T cells indicate that direct binding alone may not suffice for robust inhibition of TTL activity in cellular contexts. Instead, effective inhibition appears to require more subtle structural modulations that lead to impaired interaction of TTL with the α/β-tubulin heterodimer.

## Conclusion

In summary, we report herein the discovery and characterization of **CEM198**, a pyrimido[4,5-*d*]pyrimidine derivative, as a potent and selective antiproliferative agent. Although originally designed as a potential kinase inhibitor, comprehensive chemoproteomic profiling revealed that **CEM198** does not bind any kinase within the kinome coverage of the kinobeads platform. Instead, **CEM198** engages TTL with high affinity, representing the first small-molecule TTL ligand. Subsequent cellular target engagement studies demonstrated that **CEM198** and selected analogs inhibit TTL-mediated tyrosination of α-tubulin in a cell-type-dependent manner. Intriguingly, both **CEM198** and its analog **12** also act as tubulin polymerization inhibitors, inducing microtubule destabilization and mitotic arrest. Comparative analyses with well-established microtubule destabilizers revealed that the biological effects of **CEM198** extend beyond canonical TTL inhibition and likely arise from a direct binding to TTL combined with modulation of α/β-tubulin conformational states that restrict their productive interaction with TTL.

Altogether, our study identifies a previously unrecognized chemical scaffold capable of simultaneously perturbing the tubulin tyrosination–detyrosination cycle and microtubule polymerization. These findings not only provide valuable chemical probes for dissecting the biology of tubulin post-translational modifications but also highlight the potential of **CEM198**-derived molecules as leads for the development of anticancer agents targeting both microtubule dynamics and tubulin-modifying enzymes.

## Acknowledgements

A. Roll-Mecak group for TTL plasmid. Supported by the Deutsche Forschungsgemeinschaft (DFG, German Research Foundation) with funds from SFB1309 (Chemical Biology of Epigenetic Modifications), project 401883058 (S. L.) and 325871075 (P. K.)

