## Supplementary Files for "Chemoproteomics identifies a pyrimido[4,5-*d*]pyrimidine analogue as a tubulin-tyrosine ligase binder"

### Figures

| | 72 hours treatment | | | | | | | | | | | 24 hours treatment | |
|---|---|---|---|---|---|---|---|---|---|---|---|---|---|
| | **Capan-1** | **HCT-116** | **LN-229** | **NCI-H460** | **SH-SY5Y** | **Molt-4** | **HL-60** | **K-562** | **Z-138** | **Hek293T** | **PBMC** | **SH-SY5Y** | **Hek293T** |
| **CEM198** | 2.1 ± 0.5 | 3.6 ± 4.6 | 1.1 ± 1.0 | 0.6 ± 0.2 | 5.0 ± 1.9 | 3.4 ± 0.3 | 1.0 ± 0.8 | 0.6 ± 0.3 | 1.4 ± 1.3 | 4.7 ± 2.4 | >25 | >50 | >50 |
| **11** | 5.3 ± 2.0 | 17.2 ± 8.3 | 20.6 ± 5.1 | 24.6 ± 15.0 | 11.2 ± 5.9 | 3.5 ± 0.5 | 2.4 ± 1.3 | 4.9 ± 2.0 | 3.3 ± 1.0 | 13.6 ± 6.5 | >25 | >50 | >50 |
| **12** | 3.5 ± 4.2 | 5.5 ± 1.6 | 4.5 ± 1.5 | 5.6 ± 2.6 | 2.4 ± 1.1 | 1.8 ± 0.8 | 3.0 ± 1.0 | 3.2 ± 2.6 | 1.9 ± 1.2 | 3.1 ± 1.1 | >25 | >50 | >50 |

All values are mean ± S.D.

**Table S1.** (IC50 (µM) values for nine cancer cell lines, Hek293T, and PBMCs after 72 hours of treatment, and for SH-SY5Y and Hek293T after 24 hours of treatment

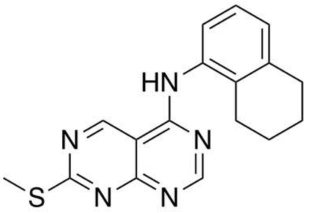

**5**

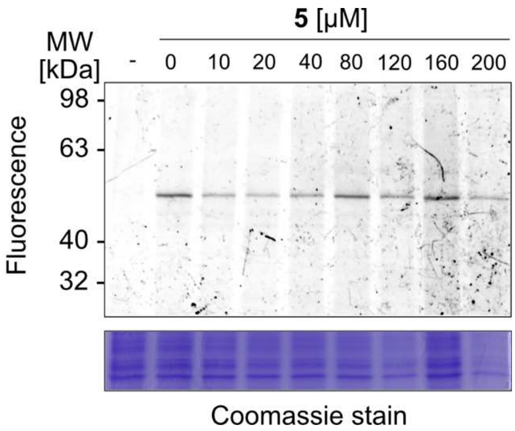

**Figure S1.** TTL activity assay in SH-SY5Y cells

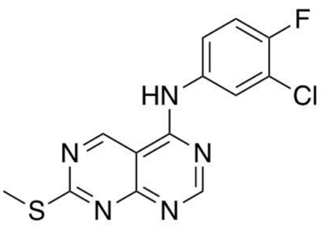

**6**

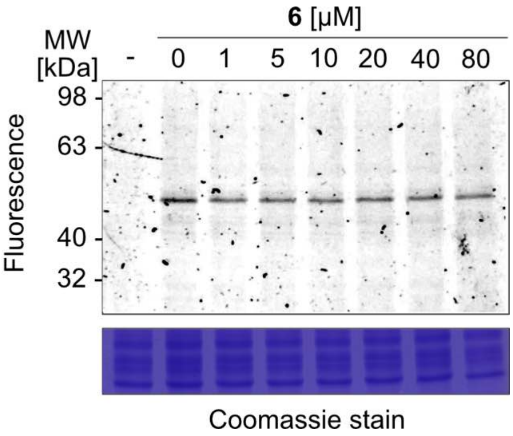

**Figure S2.** TTL activity assay in SH-SY5Y cells.

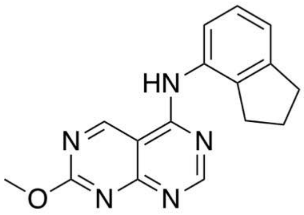

**7**

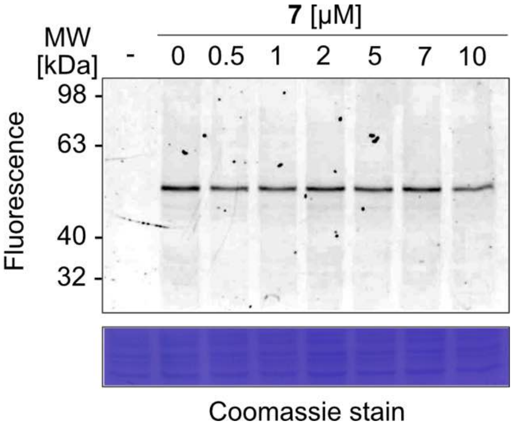

**Figure S3.** TTL activity assay in SH-SY5Y cells.

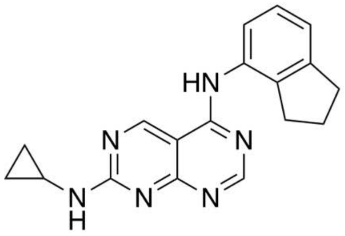

**8**

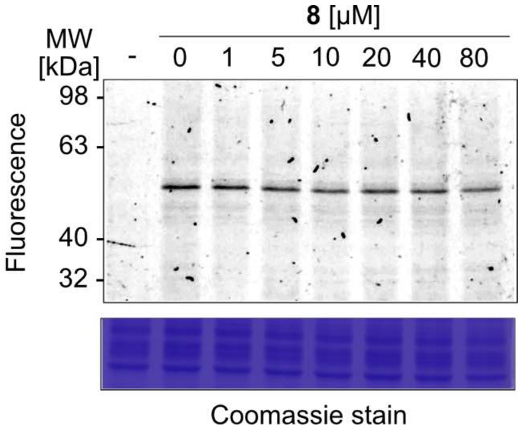

**Figure S4.** TTL activity assay in SH-SY5Y cells.

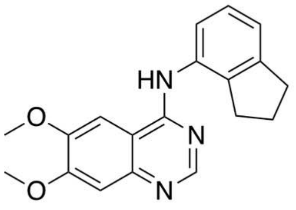

**9**

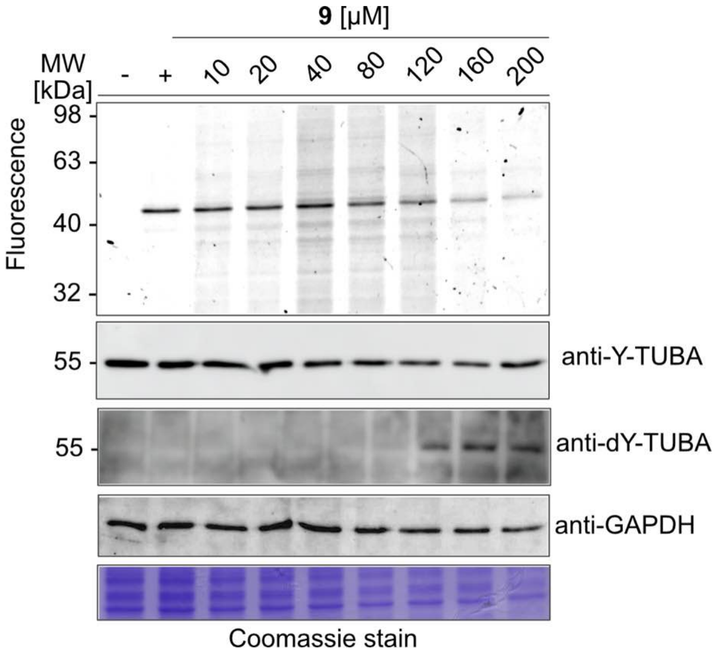

**Figure S5.** TTL activity assay in HEK293T cells.

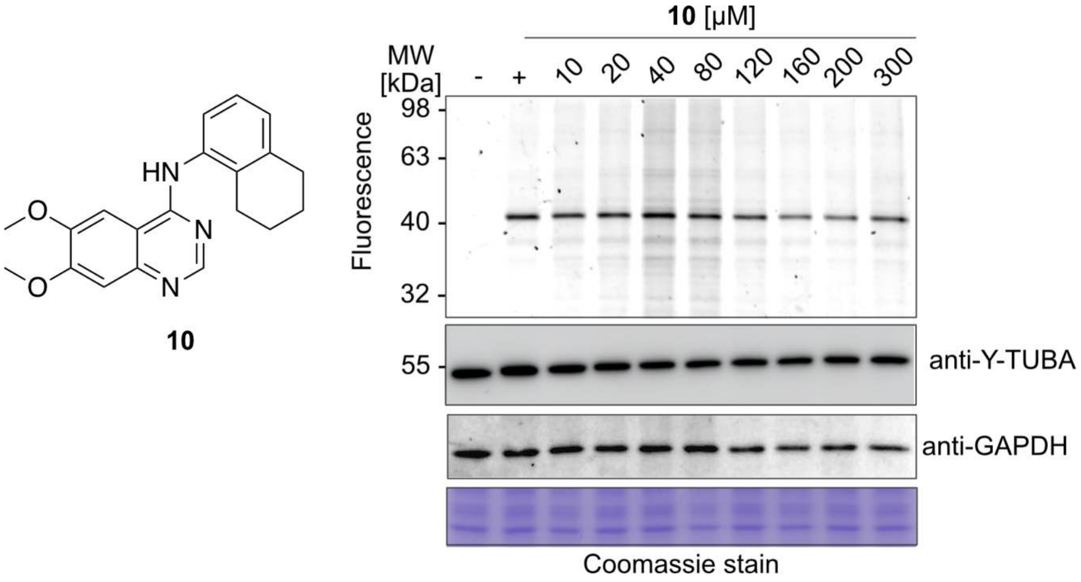

**Figure S6.** TTL activity assay in HEK293T cells.

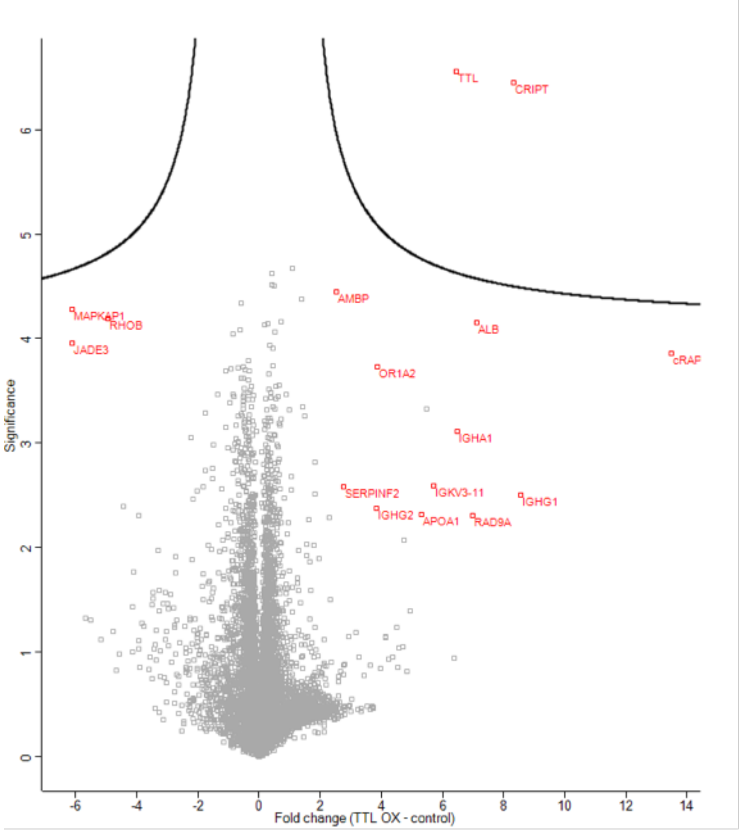

**Figure S7.** Volcano plot showing comparison of HEK293T cells with and without TTL overexpression; $n = 4$, significance is in -$\log_{10}$($p$-value), fold change is difference between conditions in label-free quantification (LFQ) intensity in $\log_2$ scale.

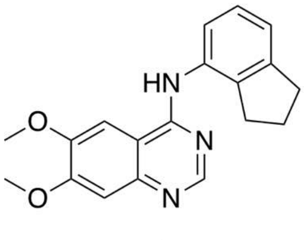

**9**

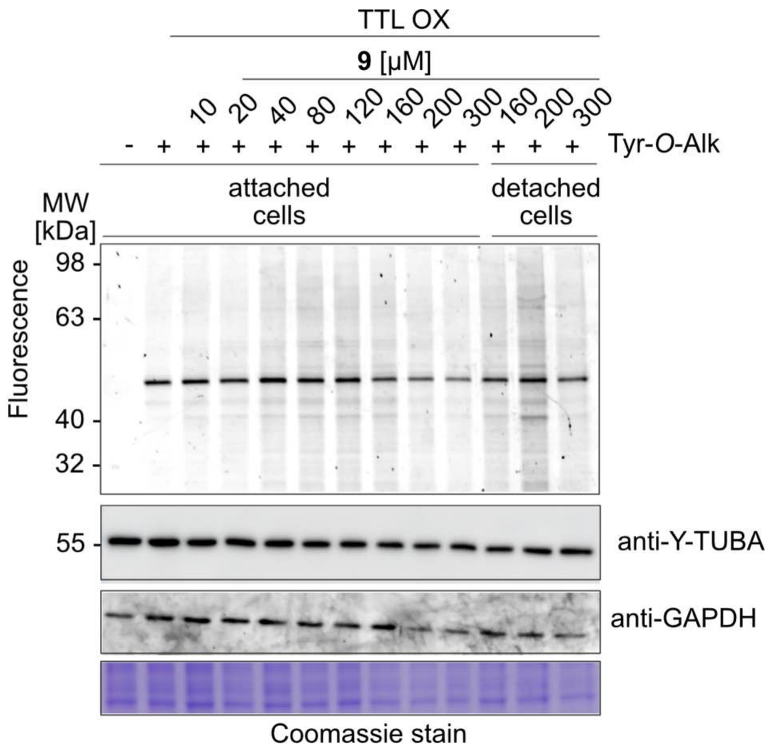

**Figure S8.** TTL activity assay in HEK293T cells overexpressing TTL.

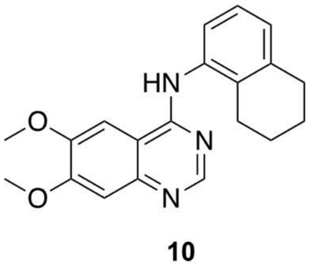

**10**

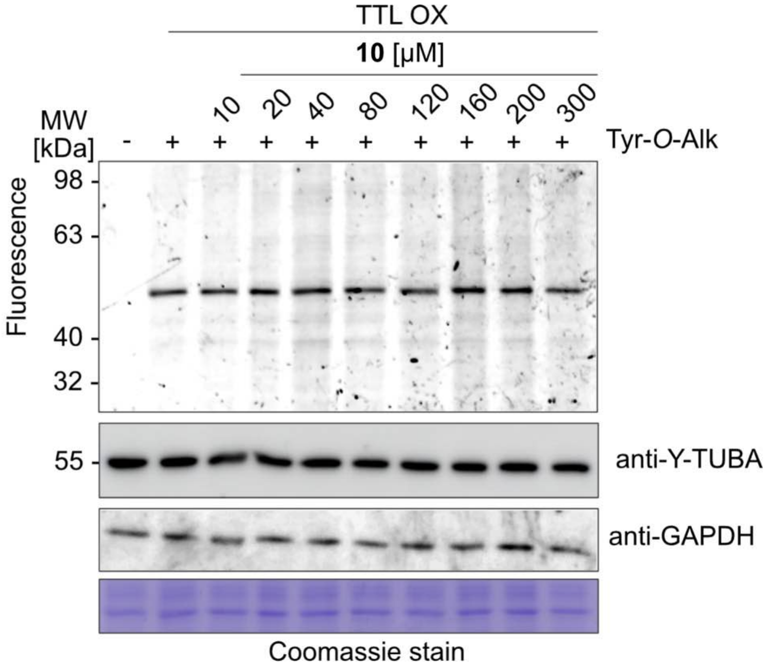

**Figure S9.** TTL activity assay in HEK293T cells with TTL OX.

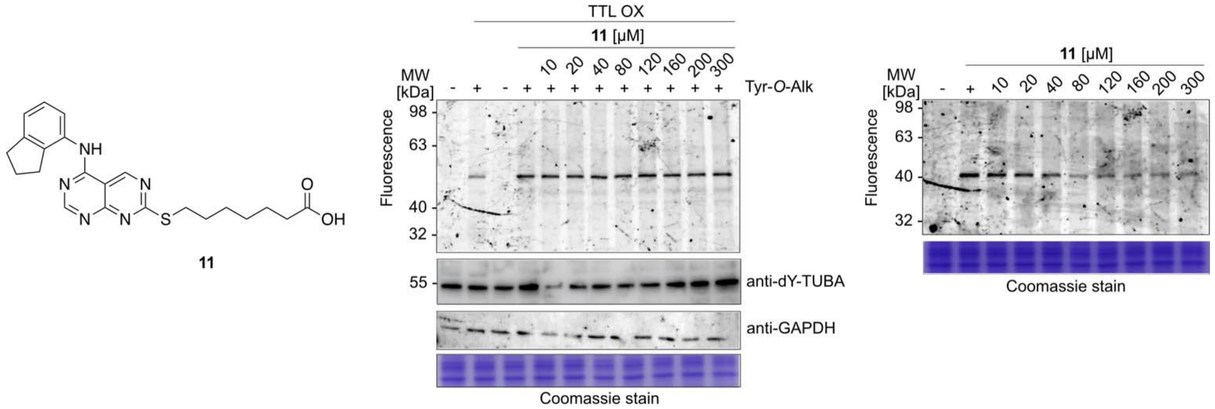

**Figure S10.** TTL activity assay in HEK293T cells with TTL OX.

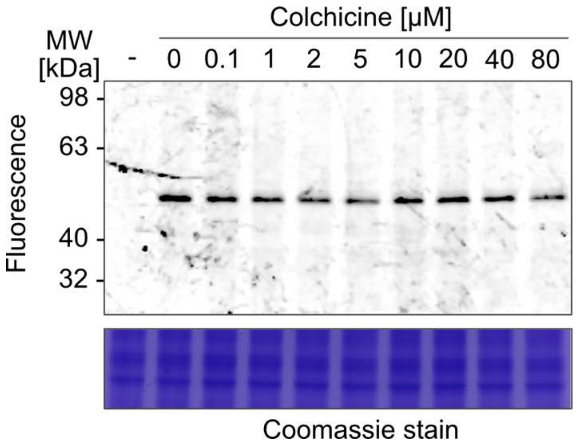

**Figure S11.** Colchicine activity in TTL assay.

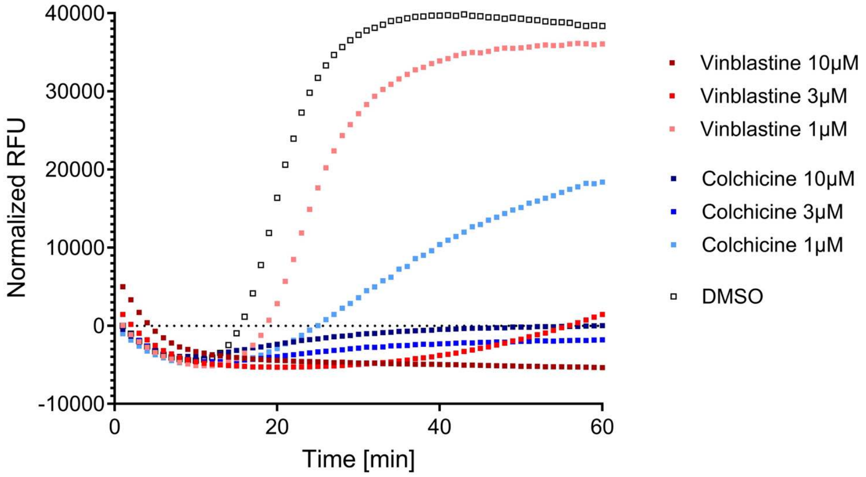

**Figure S12**. Colchicine and vinblastine in tubulin polymerization assay

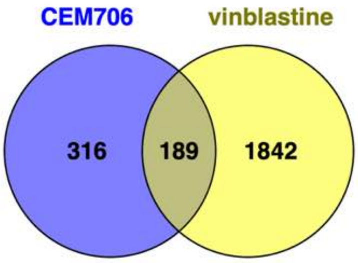

**Figure S13.** Venn diagram showing the overlap of all dysregulated proteins in **12** (CEM706) and vinblastine treated HEK293T cells.

#### Experimental Part

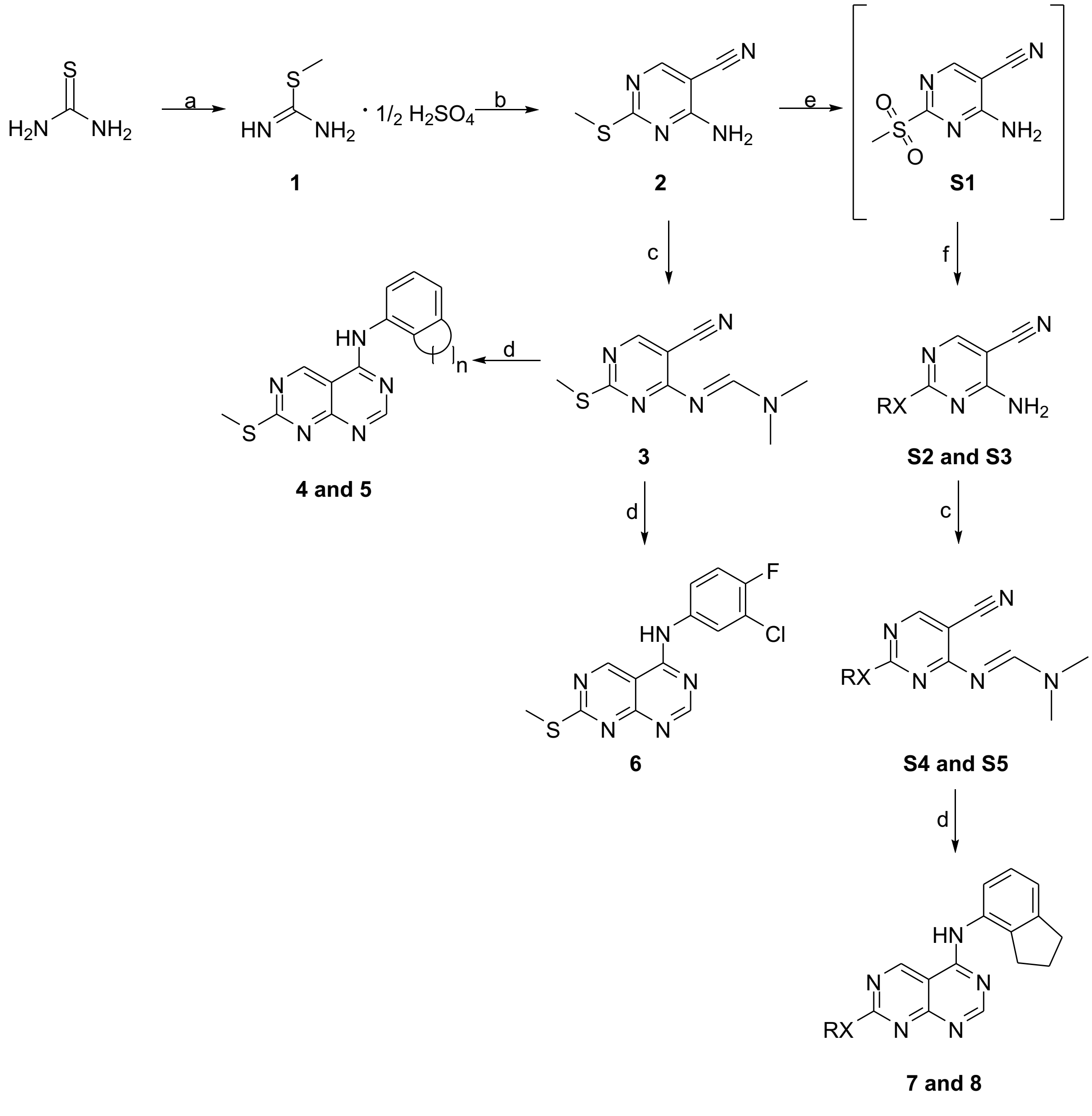

**Scheme S1.** Reagents and conditions: (a) dimethyl sulfate, $H_2O$, 80 °C; (b) 2-(ethoxymethylene)malononitrile, triethylamine, ethanol abs., rt; (c) DMF-DMA, toluene, reflux; (f) appropriate aniline, $CH_3CO_2H$, reflux; (e) *m*CPBA, $CH_2Cl_2$, 25 °C; (d) cyclopropylamine, THF, 25 °C or MeONa, MeOH, 25 °C.

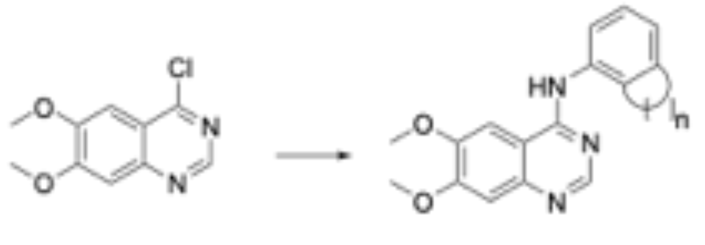

**9 and 10**

**Scheme S2.** Reagents and conditions: indan-4-amine or 5,6,7,8-tetrahydronaphthalen-1-amine, ethanol, 130 °C, autoclave.

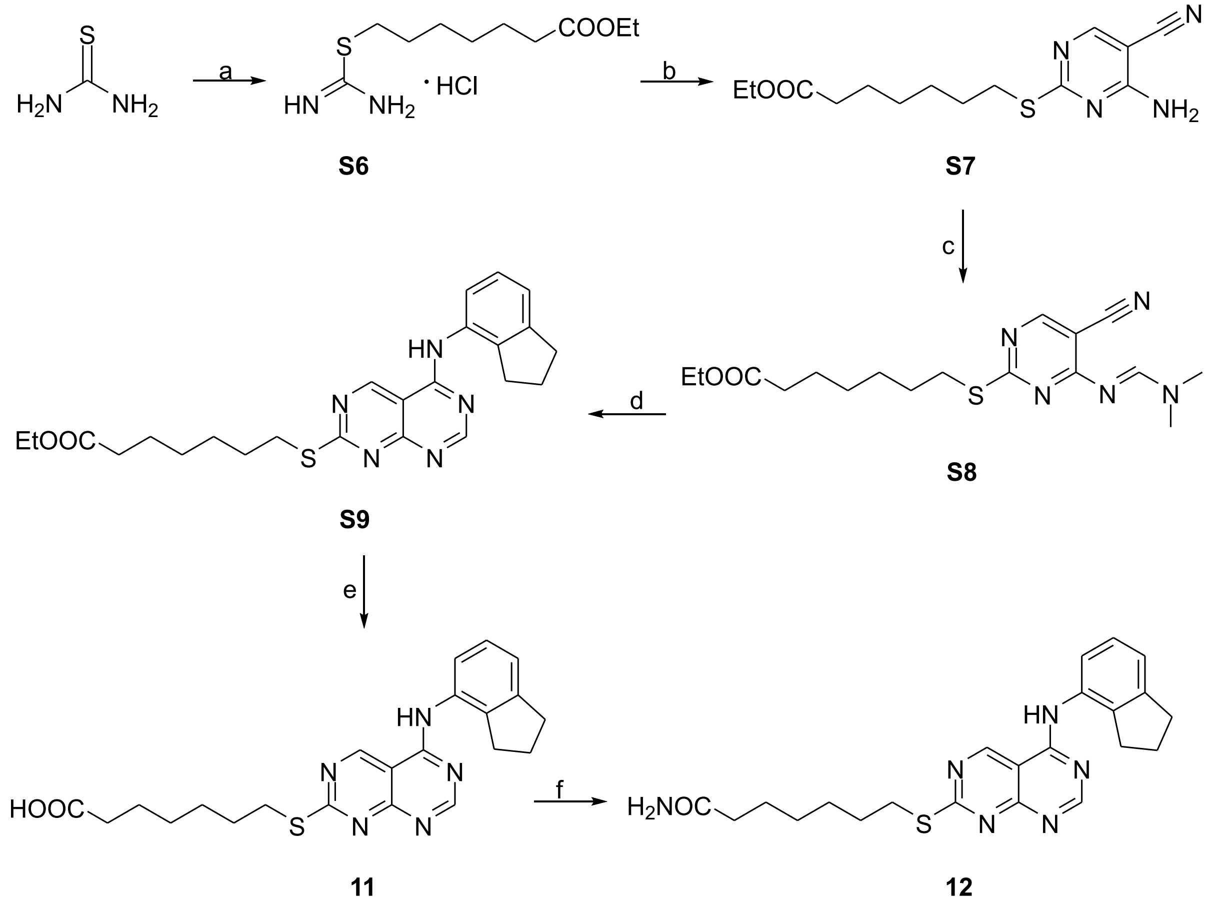

**Scheme S3.** Reagents and conditions: (a) Ethyl 6-bromoethanoate, ethanol abs., reflux; (b) 2-(ethoxymethylene)malononitrile, triethylamine, ethanol abs., rt; (c) DMF-DMA, toluene, reflux; (d) indan-4-amine, $CH_3CO_2H$, reflux; (e) NaOH, EtOH, 25 °C; (f) CDI, $NH_3$, MeOH, 25 °C.

*General Information*

All commercially available reagents and solvents were obtained from Alfa Aesar (Ward Hill, MA, USA) and used as received without further purification. Melting points were determined using a Büchi apparatus and are reported as uncorrected values. One-dimensional ($^1$H NMR, $^{13}$C NMR) and two-dimensional (COSY, NOESY, HMBC, HSQC-DEPT135) spectra were recorded on a Bruker Avance III-600 MHz spectrometer (Karlsruhe, Germany). Chemical shifts (*δ*) are reported in ppm and coupling constants (*J*) are in Hz. Signal multiplicities are designated as follows: s (singlet), d (doublet), t (triplet), q (quartet), dd (doublet of doublets), and m (multiplet). Reaction progress was monitored by analytical thin-layer chromatography on pre-coated silica gel 60 F254 plates (Merck, 0.25 mm). Mass spectrometric analyses were performed using a UPLC-Triple TOF-MS system (UPLC: Acquity, Waters, Milford, MA, USA; MS: SCIEX Triple TOF 5600+, Framingham, MA, USA).

**Methyl carbamimidothioate sulfate.** Dimethyl sulfate (13.80 g, 110.00 mmol) was added dropwise to a suspension of thiourea (15.20 g, 200.00 mmol) in water (7 mL) and the reaction mixture was heated under reflux for 12 hrs. After completion of the reaction, the mixture was cooled to room temperature, ethanol 95% (20 mL) was added and the solution was cooled to 0 °C and filtered. The filter cake was washed with cold ethanol (10 mL), while the white solid was dried over phosphorus pentoxide to afford the title compound (23 g), which was used for the next step without further purification. Yield = 82 %. $^{1}$H NMR and $^{13}$C NMR are in agreement with the reported data.[1]

**4-Amino-2-(methylthio)pyrimidine-5-carbonitrile (1).** To a solution of methyl carbamimidothioate sulfate (10 g, 3.6 mol) in abs. ethanol (65 mL), 2-(ethoxymethylene)malononitrile (8.79 g, 7.2 mol) and triethylamine (12 mL, 8.64 mol) were added. The resulting mixture was stirred at room temperature for 2 h. After completion of the reaction, the precipitate was filtered and washed with cold ethanol to afford the title compound **1** (5.4 g). Yield = 89 %. $^{1}$H NMR and $^{13}$C NMR are in agreement with the reported data.[1]

***N*-(2,3-dihydro-1*H*-inden-4-yl)-7-(methylthio)pyrimido[4,5-*d*]pyrimidin-4-amine (Cem198, 4).** To a suspension of compound **1** (166 mg, 1 mmol) in dry toluene (20 mL), N,N-dimethylformamide-dimethylacetal (133 µL, 1 mmol) was added, and the resulting mixture was stirred at 110 °C for 6 hrs. After completion of the reaction, the resulting mixture was vacuum evaporated. The oily residue was purified by column chromatography (silica gel, $CH_2$ $Cl_2$/ MeOH: 50/1 - 20/1) to afford **3** as a mixture of E/Z isomers that was added without any further purification to a solution of indan-4-amine (133 mg, 1 mmol) in glacial acetic acid (2 mL). The resulting mixture was stirred under reflux for 2.5 h and evaporated to dryness. The oily residue was purified by column chromatography (silica gel, $CH_2Cl_2$ / MeOH: 20 / 1.15 - 12 / 1 to afford **Cem198** (62 mg, 20%). $^{1}$H NMR (600 MHz, DMSO-*d*$_6$) δ (ppm) 10.45 (s, 1H), 9.68 (s, 1H), 8.60 (s, 1H), 7.19 (m, 3H), 2.93 (t, *J* = 7.5 Hz, 2H), 2.75 (t, *J* = 7.5 Hz, 2H), 2.60 (m, 3H), 1.98 (t, *J* = 7.6 Hz, 2H). $^{13}$C NMR (151 MHz, DMSO-*d*$_6$) δ 175.6, 162.7, 162.0, 159.3, 157.7, 145.4, 140.0, 133.5, 126.7, 123.5, 122.5, 104.3, 32.7, 31.1, 24.6, 13.8. ESI – HRMS calcd for $C_{16}H_{14}N_5S^-$: [M - H]$^-$ 308.0975 found 308.0969.

**7-(methylthio)-*N*-(5,6,7,8-tetrahydronaphthalen-1-yl)pyrimido[4,5-*d*]pyrimidin-4-amine (5).** Compound **5** was prepared using an analogous procedure to that of **Cem198**, with 5,6,7,8-

tetrahydronaphthalen-1-amine as the starting material. Yield: 30 %. Mp.: 285.0°C ($CH_2Cl_2$). $^1$H NMR (600 MHz, $CD_3OD$ / CDCl3 3/1) δ (ppm) 9.49 (s, 1H), 8.51 (s, 1H), 7.18 (t, *J* = 7.0 Hz, 1H), 7.12 (m, 2H), 2.85 (t, *J* = 6.0 Hz, 2H), 2.67 (s, 5H), 1.85 – 1.75 (m, 4H). $^{13}$C NMR (151 MHz, $CD_3OD$) δ (ppm) 178.6, 163.6, 163.2, 162.1, 157.9, 140.0, 136.5, 135.6, 129.9, 127.0, 125.7, 105.4, 30.6, 25.9, 23.8, 23.8, 14.5. ESI – HRMS calcd for $C_{17}H_{16}N_5S^-$: [M - H]$^-$ 322.1132 found 322.1139.

***N*-(3-chloro-4-fluorophenyl)-7-(methylthio)pyrimido[4,5-*d*]pyrimidin-4-amine (6).** Compound **6** was prepared using an analogous procedure to that of **Cem198**, using 3-chloro-4-fluoroaniline as the starting material. Yield: 46%. Mp.: >300 °C (EtOAc); $^1$H NMR (600 MHz, DMSO-*d6*) δ (ppm) 10.50 (s, 1H), 9.75 (s, 1H), 8.81 (s, $D_2O$ exchang., 0.5H), 8.14 (dd, *J* = 19.6 Hz, 16.4 Hz, 1H), 7.77 (m, 1H), 7.50 (t, *J* = 9.0 Hz, 1H), 2.62 (s, 3H); $^{13}$C NMR (51 MHz, DMSO-*d6*) δ (ppm) 175.8, 162.3, 161.8, 161.4, 160.5, 158.6, 157.5, 156.4, 151.5, 124.1, 122.9, 122.8, 119.2, 118.8, 117.0, 116.6, 115.6, 104.5, 13.8, 13.4. ESI – HRMS calcd for $C_{13}H_{10}ClFN_5S^+$ [M + H]$^+$ 322.0324 found 322.0330.

**2-Amino-4-(methylsulfonyl)benzonitrile (S1).** A suspension of 4-amino-2-(methylthio)pyrimidine-5-carbonitrile (1.0 g, 6.02 mmol, **2**) and m-chloroperbenzoic acid (3.2 g, 19.82 mmol) in dry $CH_2Cl_2$ (80 mL), was stirred at room temperature for 5 hrs. After completion of the reaction, THF was added (50 mL), and the resulting precipitate was filtered and washed with THF to provide the title compound **S1** (703 mg). Yield = 59%. $^1$H NMR and $^{13}$C NMR are in agreement with the reported data.[1]

**4-amino-2-methoxypyrimidine-5-carbonitrile (S2).** To a suspension of 2-amino-4-(methylsulfonyl)benzonitrile (98 mg, 0.50 mmol, **S1**) in dry methanol (10 mL), sodium methoxide (28 mg, 1.50 mmol) was added, and the resulting mixture was stirred at room temperature for 2 h. After completion of the reaction, the solution was evaporated to dryness and the extracted with EtOAc / water to afford the title compound **S2** (45 mg). Yield: 60%). M.p.: 221 °C (EtOH). $^1$H NMR and $^{13}$C NMR are in agreement with the reported data.[2]

**4-Amino-2-(cyclopropylamino)pyrimidine-5-carbonitrile (S3).** To a suspension of 2-amino-4-(methylsulfonyl)benzonitrile (784 mg, 4 mmol, **S1**) in dry THF (10 mL), cyclopropylamine (228 mg, 4 mmol) was added and the resulting mixture was stirred at room temperature for 14 h. After completion of the reaction, the solution was evaporated to dryness and the title compound **S3** was purified by column chromatography (silica gel, $CH_2Cl_2$/ MeOH: 50/1 - 20/1) to afford 462 mg of the title compound. Yield = 66%. $^1$H NMR and $^{13}$C NMR are in agreement with the reported data.[1]

**N-(2,3-dihydro-1*H*-inden-4-yl)-7-methoxypyrimido[4,5-*d*]pyrimidin-4-amine (7).** Compound **7** was prepared using an analogous procedure to that of **Cem198**, using 4-amino-2-methoxypyrimidine-5-carbonitrile (**S2**) and indan-4-amine, as starting materials. Yield: 40 %. M.p.: 267-268 °C (EtOAc). $^1$H NMR: (600 MHz, DMSO-*d6*) δ (ppm) 10.36 (s, 1H), 9.80 (s, 1H),

8.61 (s, $D_2O$ exchang., 1H), 7.23 – 7.17 (m, 3H), 4.04 (s, 3H), 2.95 (t, J = 7.4 Hz, 2H), 2.77 (t, J = 7.4 Hz, 2H), 2.00 (t, J = 7.5 Hz, 2H). $^{13}C$ NMR (151 MHz, DMSO-*d6*) δ 166.8, 164.1, 162.7, 161.3, 159.2, 145.3, 140.0, 133.6, 126.7, 123.5, 122.4, 103.5, 55.1, 32.7, 31.0, 24.6. ESI – HRMS calcd for $C_{16}H_{14}N_5O^-$[MH$^-$] 292.1204 found 292.1202.

***N*$^2$-cyclopropyl-*N*$^5$-(2,3-dihydro-1*H*-inden-4-yl)pyrimido[4,5-*d*]pyrimidine-2,5-diamine** (**8**). Compound **8** was prepared using an analogous procedure to that of **Cem198**, using 4-amino-2-(cyclopropylamino)pyrimidine-5-carbonitrile (**S3**) and indan-4-amine, as starting materials. $^1H$ NMR and $^{13}C$ NMR are in agreement with the reported data.[1]

***N*-(2,3-dihydro-1*H*-inden-4-yl)-6,7-dimethoxyquinazolin-4-amine (9).** A suspension of 4-chloro-6,7-dimethoxyquinazoline (224 mg, 1 mmol) and indan-4-amine (133 mg, 1 mmol) in absolute ethanol (10 mL) was stirred at 130 °C in an autoclave aparatus for 3 h. After completion of the reaction, ethanol was vacuum evaporated, and the resulting oily residue was dissolved in water and basified with 5 % $Na_2CO_3$ solution (pH~10). The mixture was washed with EtOAc (3 x 50 mL), and the combined organic layers were dried over anhydrous $Na_2SO_4$ anc concentrated to dryness. The residue was purified by column chromatography (silica gel, EtOAc) to afford compound **16** (289 mg, 90%). $^1H$ NMR and $^{13}C$ NMR are in agreement with the reported data.[3]

**6,7-dimethoxy-*N*-(5,6,7,8-tetrahydronaphthalen-1-yl)quinazolin-4-amine (10).** Compound **10** was prepared using an analogous procedure to that of compound **9**, using 5,6,7,8-tetrahydronaphthalen-1-amine, as starting material. Yield = 92 %. $^1H$ NMR and $^{13}C$ NMR are in agreement with the reported data.[3]

**ethyl 7-(carbamimidoylthio)heptanoate hydrobromide (S6).** A suspension of thiourea (1.5 g, 20 mmol) and ethyl 6-bromoethanoate (4 mL, 20 mmol) in absolute ethanol (10 mL) was stirred under reflux for 12 h. After completion of the reaction, ethanol was vacuum evaporated to afford compound **S6**, practically pure, which was used for the next step without any further purification. $^1H$ NMR (600 MHz, $CDCl_3$) δ (ppm) 8.80 (brs, 2H), 8.40 (brs, 2H), 4.13 (q, *J* = 7.2 Hz, 2H), 3.31 (t, *J* = 7.4 Hz, 2H), 2.32 (t, *J* = 7.6 Hz, 2H), 1.74 (m, 2H), 1.62 (m, 2H), 1.49 (m, 2H), 1.37 (m, 2H), 1.27 (t, *J* = 7.3 Hz, 3H). $^{13}C$ NMR (151 MHz, $CDCl_3$) δ (ppm) 174.4, 172.0, 60.6, 34.2, 31.7, 28.4, 28.2, 28.1, 24.6, 14.3.

**ethyl 7-((4-amino-5-cyanopyrimidin-2-yl)thio)heptanoate (S7.** Compound **S7** was prepared using an analogous procedure to that of compound **2**, using ethyl 7-(carbamimidoylthio)heptanoate hydrochloride (**S6**) and 2-(ethoxy methylene)malononitrile, as starting materials. Yield: 81 %). $^1H$ NMR (600 MHz, $CDCl_3$) δ (ppm) 8.30 (s, 1H), 5.64 (s, 2H), 4.16-4.11 (m 2H), 3.10 – 3.03 (m, 2H), 2.31 (m, 2H), 1.76 – 1.69 (m, 2H), 1.68-1.63 (m, 2H), 1.49-1.43 (m, 2H), 1.42 – 1.37 (m, 2H), 1.26 (t, *J* = 7.1 Hz, 3H). $^{13}C$ NMR (151 MHz, $CDCl_3$) δ (ppm) 174.2, 161.7, 160.6, 158.7, 114.8, 86.5, 60.5, 34.4, 30.9, 28.8, 28.3, 28.3, 24.7, 14.4.

**ethyl 7-((5-((2,3-dihydro-1*H*-inden-4-yl)amino)pyrimido[4,5-*d*]pyrimidin-2-yl)thio)heptanoate (S9).** Compound **S9** was prepared using an analogous procedure to that of compound **Cem198**, using ethyl 7-((4-amino-5-cyanopyrimidin-2-yl)thio)heptanoate (**S7**) and indan-4-amine, as starting materials. Yield: 62 %). $^{1}$H NMR (600 MHz, $CDCl_3$) δ (ppm) 9.17 (s, 1H), 8.79 (d, *J* = 1.0 Hz, 1H), 8.13 (s, 1H), 7.45 (d, *J* = 7.8 Hz, 1H), 7.22 (t, *J* = 7.6 Hz, 1H), 7.18 (d, *J* = 7.5 Hz, 1H), 4.11 (q, *J* = 7.1 Hz, 2H), 3.26 (t, *J* = 7.3 Hz, 2H), 3.00 (t, *J* = 7.5 Hz, 2H), 2.85 (t, *J* = 7.4 Hz, 2H), 2.28 (t, *J* = 7.5 Hz, 2H), 2.11 (p, *J* = 7.5 Hz, 2H), 1.76 (p, *J* = 7.4 Hz, 2H), 1.62 (p, *J* = 7.6 Hz, 2H), 1.47 (p, *J* = 7.4 Hz, 2H), 1.38 – 1.31 (m, 2H), 1.24 (dd, *J* = 7.7, 6.6 Hz, 3H). $^{13}$C NMR (151 MHz, $CDCl_3$) δ (ppm) 177.6, 173.9, 163.22, 162.7, 159.6, 155.4, 146.6, 138.8, 133.2, 127.7, 123.4, 122.3, 104.3, 60.4, 34.5, 33.4, 31.4, 31.1, 28.8, 28.7, 28.6, 25.1, 24.9, 14.4. ESI – HRMS calcd for $C_{24}H_{28}N_5O_2S^-$ [M - H]$^-$ 450.1969 found 450.1973

**7-((5-((2,3-dihydro-1*H*-inden-4-yl)amino)pyrimido[4,5-*d*]pyrimidin-2-yl)thio)heptanoic acid (11).** To a suspension of compound **S9** (100.0 mg, 0.22 mmol) in ethanol (5 mL) was added dropwise a cold 40% NaOH solution (100 µL). The mixture was stirred for 12 hrs at room temperature and then pured into water and acidified with 9% HCl solution (pH ~ 5). The resulting precipitate was filtered and air dried to afford 65 mg of the title compound, practically pure. Yield: 70%. Mp.: 181.8 °C ($CH_3COOH$/EtOAc). $^{1}$H NMR (600 MHz, $CD_3OD$) δ (ppm) 9.47 (s, 1H), 8.60 (s, 1H), 7.20-7.18 (m, 2H), 7.13 (t, *J* = 7.3 Hz, 2H), 3.28 – 3.18 (m, 2H), 3.00 – 2.91 (m, 2H), 2.81– 2.75 (m, 2H), 2.29 – 2.24 (m, 2H), 2.08 – 2.00 (m, 2H), 1.79 – 1.71 (m, 2H), 1.63 – 1.53 (m, 2H), 1.51 – 1.41 (m, 2H), 1.37 (q, *J* = 7.5 Hz, 3H). $^{13}$C NMR (151 MHz, $CD_3OD$-CDCl3 3/1) δ 178.0, 177.2, 162.6, 162.1, 160.7, 157.6, 147.0, 141.2, 133.6, 127.8, 124.2, 124.0, 105.2, 34.6, 33.8, 32.0, 3.7, 29.4, 29.1, 25.7, 25.4. ESI – HRMS calcd for $C_{22}H_{24}N_5O_2S^-$ [M - H]$^-$ 422.1656 found 422.1659

**7-((5-((2,3-dihydro-1*H*-inden-4-yl)amino)pyrimido[4,5-*d*]pyrimidin-2-yl)thio)heptanamide (12).** To a solution of compound **11** (58 mg, 0.14 mmol) in dry DMF (5 mL) was added 1,1'-carbonyldiimidazole (28 mg, 0.16 mmol), and the resulting mixture was stirred at room temperature under argon for 4h. Subsequently, a cold 24% $NH_3$ solution (15 µL) was added, and the resulting syspension was stirred at room temperature for an additional 24 hrs. After completion of the reaction, the mixture was diluted with water (30 mL) and washed succesively with EtOAc (3 x 20 mL). The combined organic layers were dried over anhydrous $Na_2SO_4$ and concentrated under reduced pressure. The residue was purified by column chromatography (silica gel, $CH_2Cl_2$/ MeOH: 50/1 - 20/1) to afford 29 mg of the title compound. Yield: 49%. Mp.: 198.4°C (EtOAc/*n*-pentane). $^{1}$H NMR (600 MHz, $CDCl_3$) δ (ppm) 9.49 (s, 1H), 8.73 (s, 1H), 7.84 (s, 1H), 7.33 (dd, *J* = 7.4, 1.5 Hz, 1H), 7.21 – 7.15 (m, 2H), 7.13 (s, 2H), 3.23 – 3.16 (m, 2H), 2.97 (t, *J* = 7.5 Hz, 2H), 2.84 (t, *J* = 7.4 Hz, 2H), 2.22 (t, *J* = 7.5 Hz, 2H), 2.10 – 2.04 (m, 3H), 1.72 (q, *J* = 7.5 Hz, 2H), 1.63 (p, *J* = 7.6 Hz, 2H), 1.46 - 1.41 (m, 2H), 1.39 – 1.34 (m, 2H). $^{13}$C NMR (151 MHz, $CDCl_3$) δ (ppm) 177.2, 176.1, 163.0, 162.6, 159.8,

156.5, 146.5, 139.6, 133.4, 127.5, 123.4, 122.8, 104.6, 35.9, 33.4, 31.4, 31.1, 28.6, 28.47, 28.37, 25.4, 25.1. ESI – HRMS calcd for $C_{22}H_{26}N_6OS^-$ [M - H]$^-$ 421.1816 found 421.1810.

#### Analytical and biochemical methods

##### Cell culture

Human cancer cell lines (Capan-1, HCT-116, NCI-H460, LN-229, SH-SY5Y, HL-60, K-562, Z-138, Molt-4) were obtained from ATCC (Manassas, VA, USA), and DND-41 cells were sourced from DSMZ (Leibniz Institute, Germany). HEK293T cells were kindly provided by Prof. J. Moffat. All media (Gibco, Life Technologies, USA) were supplemented with 10% fetal bovine serum (HyClone, GE Healthcare Life Sciences, USA). Stock solutions of test compounds were prepared in DMSO.

##### Proliferation assays

For adherent lines, Capan-1 cells were seeded at 500 cells/well, and HCT-116, NCI-H460, LN-229, and SH-SY5Y at 1,500 cells/well in 384-well plates. After overnight attachment, cells were treated with seven serial dilutions of compounds (50 µM to 3.2 nM). Suspension lines HL-60, K-562, Molt-4 and Z-138 were plated at 2,500 cells/well, and DND-41 at 5,500 cells/well under identical concentration ranges. Following 72 h incubation, viability was determined using the CellTiter 96® AQueous One Solution (MTS) assay per manufacturer's protocol. Absorbance was recorded at 490 nm using a SpectraMax Plus 384 reader, and $IC_{50}$ values were calculated from optical density data. Each compound was tested in at least two independent experiments.

##### Cell viability of normal PBMC

Peripheral blood mononuclear cells (PBMCs) were isolated from buffy coats of healthy donors (Blood Transfusion Center, Leuven, Belgium) by density gradient centrifugation using Lymphoprep (Nycomed, Oslo, Norway; density 1.077 g/mL). PBMCs were cultured in DMEM/F12 medium (Gibco) supplemented with 8% FBS and seeded at 28,000 cells/well in 384-well plates. Compounds were tested at concentrations from 25 µM to 1.6 nM. After 72 h, viability was assessed using the MTS assay. Each compound was evaluated in four independent experiments using PBMCs from different healthy donors.

##### Tubulin polymerization assay

Tubulin assembly was monitored using a fluorescence-based kit (BK011P, Cytoskeleton, Denver, CO) according to the supplier's protocol. Briefly, half-area 96-well plates were pre-warmed at 37°C for 10 min. Test and reference compounds were added as 10× stock solutions (5 µL per well, in duplicate). Polymerization buffer (2 mg/mL tubulin in 80 mM PIPES, 2 mM $MgCl_2$, 0.5 mM EGTA, pH 6.9, containing 10 µM fluorescent reporter, 15% glycerol, and 1 mM GTP) was added. Fluorescence was recorded kinetically for 60 min at 37°C using a Tecan Spark reader (Ex 350 nm, Em 435 nm; four reads per well).

**Cell Cycle Analysis by High-Content Imaging**

SH-SY5Y cells were seeded at 5,000 cells/well in 96-well clear-bottom plates. After overnight incubation, cells were treated with compounds for 24 h, fixed with 4% paraformaldehyde in PBS for 10 min, washed, and stained with 300 nM DAPI (Molecular Probes). Imaging was performed on a CX5 High Content Screening platform (Thermo Fisher Scientific) using the Cell Cycle Analysis application. At least 1,000 cells were analyzed per well.

**Preparation of kinobeads.** Kinobeads were prepared as in ref. (https://pubs.acs.org/doi/10.1021/acschembio.8b01020).

**Preparation of iCEM670.** Ethylendiamine (1 µmol, in DMSO) was reacted with DMSO-washed NHS-activated (~20 µmol/mL beads) sepharose beads (1 mL) and triethylamine (15 µL) in DMSO (2 mL) on an end-over-end shaker for 16 h at RT in the dark. (TLC with Kaiser test staining was used to monitor successful conversion). Aminoethanol (50 µL) was then added to inactivate the remaining NHS-activated carboxylic acid groups. After 2 h on an end-over-end shaker at RT, the beads were washed with DMSO (4 x 10 mL) and resuspended in anhydrous DMF (2 mL total volume). HATU (10 µmol, 100 µL of 100 mM stock in DMF), CEM670 (12 µmol, 120 µL of 100 mM stock in DMSO), Hünig's base (20 µmol, 100 µL of 200 mM stock in DMF) and triethylamine (20 µL) were then added and the beads were incubated at RT for 16 h on an end-over-end shaker. Next, the beads were washed twice with 10 mL DMF and thrice with 10 mL ethanol. Beads were stored at 4 °C in EtOH.

**Preparation of cell lysates for chemoproteomic assays.** K-562 cells were grown in IMDM medium. The medium was supplemented with 10% FBS and the cell line was tested for Mycoplasma contamination. Cells were lysed in lysis buffer (0.8% Igepal, 50 mM Tris-HCl pH 7.5, 5% glycerol, 1.5 mM $MgCl_2$, 150 mM NaCl, 1 mM $Na_3VO_4$, 25 mM NaF, 1 mM DTT and supplemented with protease inhibitors (SigmaFast, Sigma) and phosphatase inhibitors (prepared in-house according to Phosphatase inhibitor cocktail 1, 2 and 3 from Sigma-Aldrich)). The protein amount of cell lysates was determined by Bradford assay and adjusted to an Igepal concentration of 0.4% and protein concentration of 5 mg/mL.

**Chemoproteomic competition assays.** Cell lysate was pre-incubated with different doses of the CEM198 and a DMSO vehicle control for 1 h at 22 °C in an end-over-end shaker, followed by incubation with 18 µL affinity matrix (kinobeads or iCEM670) for 30 min at 30 °C in an end-over-end shaker. To assess the degree of protein depletion from lysates by the affinity matrix, a second pulldown (PDPD) with fresh beads was performed using the unbound protein fraction from the vehicle control flow through. The beads were washed (1x 1 mL of lysis buffer without inhibitors and only 0.4% Igepal, 2x 2 mL of lysis buffer without inhibitors and only 0.2% Igepal) and captured proteins were denatured with 8 M urea buffer, alkylated with 55 mM chloroacetamide and digested with Trypsin according to standard procedures. Resulting peptides were desalted on a C18 filter plate (Sep-Pak® tC18 µElution Plate, Waters), vacuum dried and stored at -20 °C until LC-MSMS measurement.

**LC-MS/MS measurement of chemoproteomic assays.** Peptides were analyzed via LC-MS/MS on a Dionex Ultimate3000 nano HPLC coupled to an Orbitrap HF-X (Thermo Fisher Scientific) mass spectrometer, operated via the Thermo Scientific Xcalibur software. Peptides were loaded on a trap column (100 µm x 2 cm, packed in house with Reprosil-Gold C18 ODS-3 5 µm resin, Dr. Maisch, Ammerbuch) and washed with 5 µL/min solvent A (0.1 % formic acid in HPLC grade water) for 10 min. Peptides were then separated on an analytical column (75 µm x 40 cm, packed in house with Reprosil-Gold C18 3 µm resin, Dr. Maisch, Ammerbuch) using a 50 min gradient ranging from 4-32 % solvent B (0.1 % formic acid, 5 % DMSO in acetonitirile) in solvent A (0.1 % formic acid, 5 % DMSO in HPLC grade water) at a flow rate of 300 nL/min.

The mass spectrometer was operated in data dependent acquisition mode, automatically switching between MS1 and MS2 spectra. MS1 spectra were acquired over a mass-to-charge (m/z) range of 360-1300 m/z at a resolution of 120,000 in the Orbitrap using a maximum injection time of 50 ms and an automatic gain control (AGC) target value of 3e6. Up to 25 peptide precursors were isolated (isolation window width of 1.3 Th, maximum injection time of 22 ms, AGC target value of 1e5), fragmented by HCD using 28 % normalized collision energy (NCE) and analyzed in the Orbitrap at a resolution of 15,000. The dynamic exclusion duration of fragmented precursor ions was set to 20 s.

**Protein identification and quantification.** Protein identification and quantification was performed using MaxQuant[2] (v 1.6.1.0) by searching the LC-MS/MS data against all canonical protein sequences as annotated in the Swissprot reference database (v03.12.15, 20193 entries, downloaded 22.03.2016) using the embedded search engine Andromeda. Carbamidomethylated cysteine was set as fixed modification and oxidation of methionine and N-terminal protein acetylation as variable modifications. Trypsin/P was specified as proteolytic enzyme and up to two missed cleavage sites were allowed. Precursor tolerance was set to 10 ppm and fragment ion tolerance to 20 ppm. The minimum length of amino acids was set to seven and all data were adjusted to 1% PSM and 1% protein FDR. Label-free quantification[2] and match between runs was enabled.

**Chemoproteomic competition assay data analysis.** For the competition assays, relative binding was calculated based on the protein intensity ratio to the DMSO control for every single inhibitor concentration. $EC_{50}$ values were derived from a four-parameter log-logistic regression with variable slope. Targets of the inhibitors were annotated manually. A protein was considered a target or interactor of a target if the resulting binding curve showed a sigmoidal curve shape with a dose dependent decrease of binding to the beads. Additionally, the number of unique peptides and MSMS counts per condition were taken into account.

**SH-SY5Y and HEK293T cell line**. The human neuroblastoma cell line SH-SY5Y (CRL-2266) or HEK293T cells were cultivated in a high-glucose Dulbecco's Modified Eagle Medium (DMEM) that was further supplemented with 10% (v/v) heat-inactivated fetal bovine serum (FBS) and 2% (v/v) L-glutamine. Cells were maintained in cell culture dishes for adherent cells at 37°C under constant humidity and 5% $CO_2$ concentration.
**Cells treatment and harvesting**. SH-SY5Y or HEK293T cells were treated with the stock solution of **Tyr-*O*-Alk** ($H_2O$ : 1M NaOH 2:1, 144 mM, sterile filtered), the final concentration in culture media was 0.3 mM. Unless otherwise stated, the cells were probe treated for 24 h. Using the same volume as for the probe-treated cells, we treated the control group with a plain solvent, the same as that used to prepare the probe stock solution.
**Lysates preparation**. To prepare a cell lysate, the cell pellet was reconstituted in 300 µL of a lysis buffer (1% NP40, 0.2% SDS in 25 mM Hepes, 7.5 pH) by sonication with an ultrasonic tip in 1 s on/ 1 s off cycles at 20% intensity for 10 s of total time. The solution was clarified by centrifugation at 4°C at 14000 rcf for 15 min. The clear supernatant was then transferred to a new 1.5 mL tube and stored at -80°C until use.
**Protein concentration measurement**. Protein concentration measurement was performed with a Pierce™ BCA Protein Assay Kit (Thermo Scientific).

**Click reaction.** The final concentration of the proteins in the samples was adjusted to 1mg/mL. Lysates were diluted to 100 µL with lysis buffer (1% NP40, 0.2% SDS in 25 mM Hepes, 7.5 pH). For each sample, 1 µL of TAMRA-$N_3$ (10 mM in DMSO), 3 µL of TCEP (100 mM in $H_2O$), and 0.125 µL TBTA (83.5 mM in DMSO) were added, vortexed, spun down, and supplemented with 2 µL of $CuSO_4$ (50 mM in $H_2O$) to initiate the reaction. The reaction mixture was incubated at RT. while shaking at 650 rpm for 1.5 h in dark.

**In-gel fluorescence analysis**. The resolution of proteins was made with the SDS-PAGE using 10% acrylamide gels. Before loading onto the gel, a protein solution (20 µL) was mixed with 5 × SDS reducing loading buffer (5 µL) (10% (w/v) SDS, 50% (v/v) glycerol, 25% (v/v) β-mercaptoethanol, 0.5% (w/v) bromphenol blue, 315 mM Tris/HCl, pH 6.8) and placed in wells. As a reference, two types of protein markers were used: BenchMark™ Fluorescent Protein Standard (Invitrogen™), Color Prestained Protein Standard, Broad Range (10-250 kDa) (New England Biolabs GmbH). Afterward, the gel was scanned on Amersham Imager 680 (GE Healthcare).
**Western blot**. After separating the proteins on a gel, they were blotted on a PVDF membrane using a Semi-Dry Blotter (Bio-Rad). Before making a blotting sandwich, a thick blot paper was soaked in a blot buffer (48 mM Tris, 39 mM glycine, 0.0375% (m/v) SDS, 20% (v/v) methanol) for 5 min, and a membrane was incubated in methanol. After the blotting, the membrane was

set in a blocking solution (0.5 g nonfat dried milk powder in 10 mL PBST (PBS + 0.5% Tween)) for 60 min to hide all nonspecific binding sites. The membrane was then placed in the primary antibody of interest solution and incubated at 4°C overnight. The next day, the membrane was washed 3 × 10 min with PBST solution before incubation with the secondary HRP-linked antibody solution at r.t. for 1 h. The membrane was then washed with PBST 3 × 10 min. Before scanning the membrane on Amersham Imager 680 (GE Healthcare), it was wetted with the ECL substrate and the peroxide solution in a 1:1 ratio. Anti-α-tubulin Antibody, tyrosinated, clone YL1/2 with catalog number MAB1864-I (Sigma-Aldrich) and anti-alpha tubulin antibody, non-tyrosinated with catalog number ABT170 (Sigma-Aldrich) were used for the analysis.

**Whole proteome samples preparation**. HEK293T cell lysates were prepared under standard conditions described above. The volume of a lysate containing 100 µg of proteins was normalized to 400 µL with lysis buffer. A mixture of hydrophilic and hydrophobic carboxylate-coated magnetic beads was washed three times with 500 µL of MS-grade $H_2O$. The sample was added to the beads and thoroughly mixed. 600 µL of absolute EtOH was added, and the mixture was incubated at r.t. for 5 min with agitation. Subsequently, the beads were washed with EtOH (80% in $H_2O$) three times. To cleave the proteins, on-beads digestion was performed. In brief, beads were reconstituted in 100 mM ammonium bicarbonate buffer alongside reducing and alkylating agent and boiled. After the samples were cooled down, trypsin was added and left overnight at 37°C. The supernatant was then placed in a new 1.5 mL tube, and the beads were washed several times with ABC buffer. After desalting the peptide mixture on a C18 column, samples were dried on a SpeedVac. The dry peptides were reconstituted in 200 µL of FA (1% in H2O) and transferred into MS vials.

**Whole proteome LC-MS/MS proteomics**. *Sample preparation and mass spectrometric measurements.* All samples were prepared in 96-well plate format using the optimized SP3 protocol according to Hughes et al. [4] Protein amount was adjusted to 20 µg in a total volume of 40 µL lysis buffer. The protein was loaded onto a mixture of hydrophilic and hydrophobic carboxylate-coated magnetic beads (Cytiva, #45152105050250 and #65152105050250, 10 µL each) prewashed three times with 100 µL of MS-grade $H_2O$ (Honeywell, #15665350). The magnetic beads with protein samples were mixed at 850 rpm, 1 min at RT. To initiate the binding, 60 µL of absolute EtOH was added, and the mixture was incubated at RT for 5 min at 850 rpm. Subsequently, the beads were washed three times with 80% (v/v) EtOH, with incubation at RT for 1 min and 850 rpm between each wash. After the last wash, the beads were resuspended in 50 µL of 100 mM ammonium acetate buffer (ABC, Sigma-Aldrich, #09689). The wash steps and ABC buffer addition was performed by a liquid handling robot (Hamilton Microlab Prep). The on-beads digestion was performed with trypsin (Promega, V5113, 1 µg) overnight 37 ˚C and 850 rpm. The resulting peptide mixture was eluted from the magnetic beads into a new 1.5 mL tube. The magnetic beads were washed with 50 and 30 µL of 1% (v/v) formic acid (TCI, # F0654) and incubated at 40 ˚C, 850 rpm for 5 min. The fractions were added to the first elution fraction. 4 µL was used for the injection. MS measurements were performed on an Orbitrap Eclipse Tribrid Mass Spectrometer (Thermo Fisher Scientific) coupled to an UltiMate 3000 Nano-HPLC (Thermo Fisher Scientific) via a nanospray Flex ion source (Thermo Fisher Scientific) equipped with column oven (Sonation) and FAIMS interface (Thermo Fisher Scientific). Peptides were loaded on an Acclaim PepMap 100 µ-precolumn cartridge (5 µm, 100 Å, 300 µm ID x 5 mm, Thermo Fisher Scientific) and separated at 40°C on a PicoTip emitter (noncoated, 15 cm, 75 µm ID, 8 µm tip, New Objective) that was in-house packed with Reprosil-Pur 120 C18-AQ material (1.9 µm, 150 Å, Dr. A. Maisch GmbH). Buffer composition: Buffer A consists of MS-grade $H_2O$ supplemented with 0.1% FA; Buffer B consists of acetonitrile supplemented with 0.1% FA. The 60-min LC gradient from 4 to 35.2 % buffer B was used. The flow rate was 0.3 µL/min.

*Data independent acquisition.* The DIC duty cycle comprised one MS1 scan followed by 30 MS2 scans. The MS2 scans had an isolation window of 4 m/z range, which overlapped with an adjacent window at the 2 m/z range.MS1 scan was conducted with Orbitrap at 60000 resolution power and a scan range of 200 – 1800 m/z with an adjusted RF lens at 30%. MS2 scans were conducted with Orbitrap at 30000 resolution power, RF lens was set to 30%. The precursor mass window was restricted to a 500 – 740 m/z range. HCD fragmentation was enabled as an activation method with a fixed collision energy of 35%. FAIMS was performed with one CV at -45V for both MS1 and MS2 scans during the duty cycle.

**Quantification and statistical analysis**. Raw files were converted in the first step with "MSConvertGUI" as a part of the "ProteoWizard" software package (http://www.proteowizard.org/download.html) to an output mzML format applying the "peakPicking" filter with "vendor msLevel=1", and the "Demultiplex" filter with parameters "Overlap Only" and "mass error" set to 10 ppm. Standalone DIA-NN software under version 1.8.1 was used for protein identification and quantification. First, a spectral library was predicted *in silico* by the software's deep learning-based spectra, RTs and IMs prediction using Uniprot *H. sapiens* decoyed FASTA (canonical and isoforms – May 2022). FASTA digest for library-free search/library generation option was enabled for this. Spectrallibrary prediction was performed in 4 batches of 10 samples each to decrease the computational load. Second, all samples (40) were processed together without spectral library generation, with a match between runs (MBR) option and precursor FDR level set at 1%. Previously generated spectral libraries were implemented during the search by presenting the command ('--lib [file name]') into the command box. DIA-NN search settings: Library generation was set to smart profiling, Quantification strategy - Robust LC. The mass accuracy and the scan window were set to 0 to allow the software to identify optimal conditions. The precursor m/z range was changed to 500-740 m/z to fit the measuring parameters. Carbamidomethylation was set as a fixed modification, oxidation of methionine and N-term acetylation were set as variable modifications. On the contrary, the small-scale samples of the 96- well plate were calculated without carbamidomethylation as a fixed modification. Statistical analysis of the DIA-NN result table “report.pg_matrix.csv” was done with Perseus 1.6.10.43. First, potential contaminants, as well as reverse peptides, were removed from the table. Then the LFQ intensities were log2-transformed. Afterward, the rows corresponding to a time point were divided into two groups – Control and Probe-treated sample. Subsequently, the groups were filtered for at least three valid values out of four rows in at least one group, and the missing values were replaced from a normal distribution with a downshift of 1.8. The $-\log_{10}(p\text{-values})$ were obtained by a two-sided one-sample Student’s t-test over replicates with the initial significance level of $p = 0.05$. Fold change values, as well as p-values, were obtained for each time point.[5, 6]

#### NMR spectra

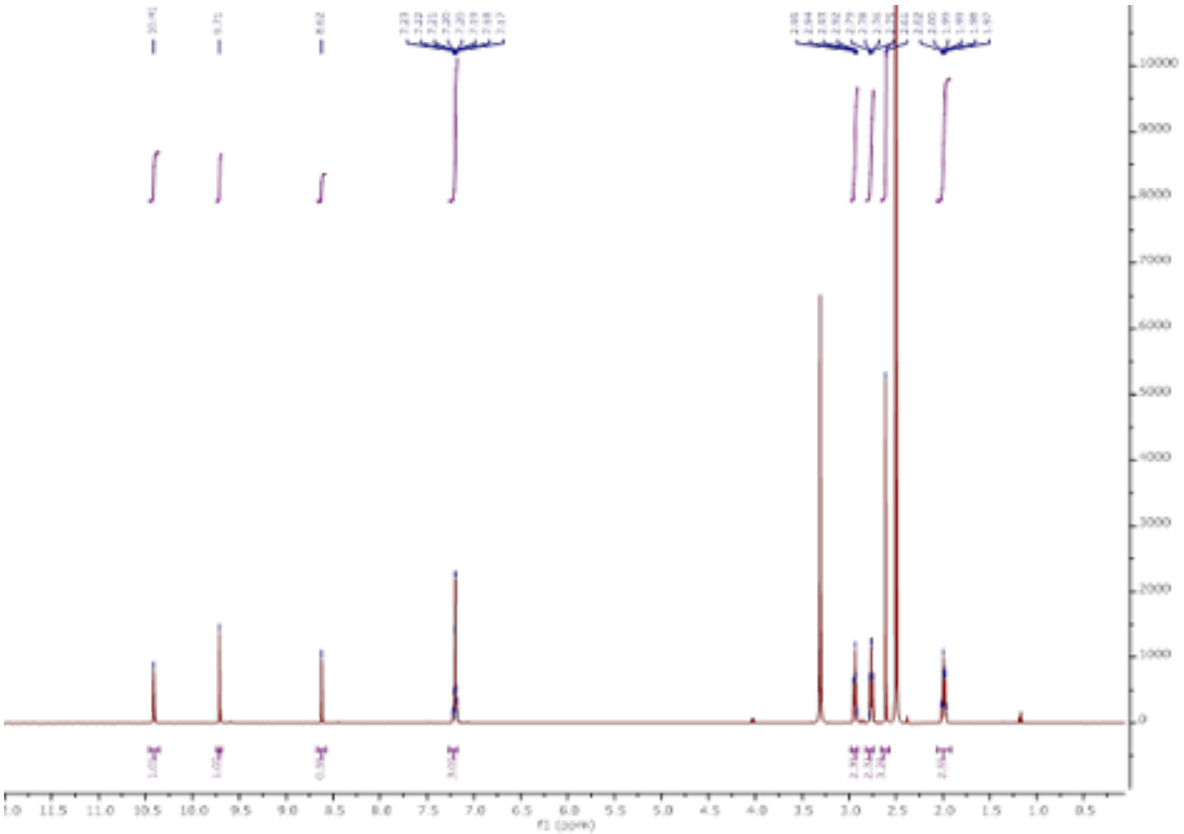

[1]H NMR spectrum of **CEM198**

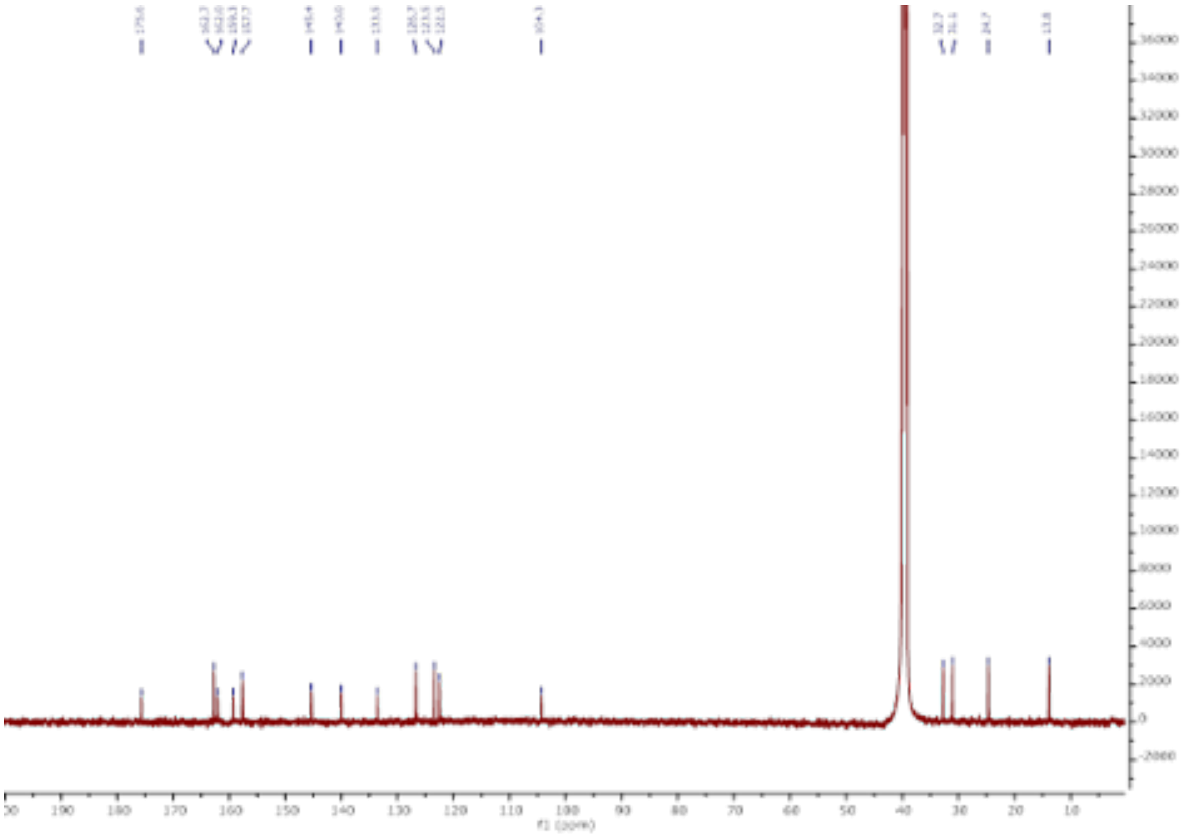

[13]C NMR spectrum of **CEM198**

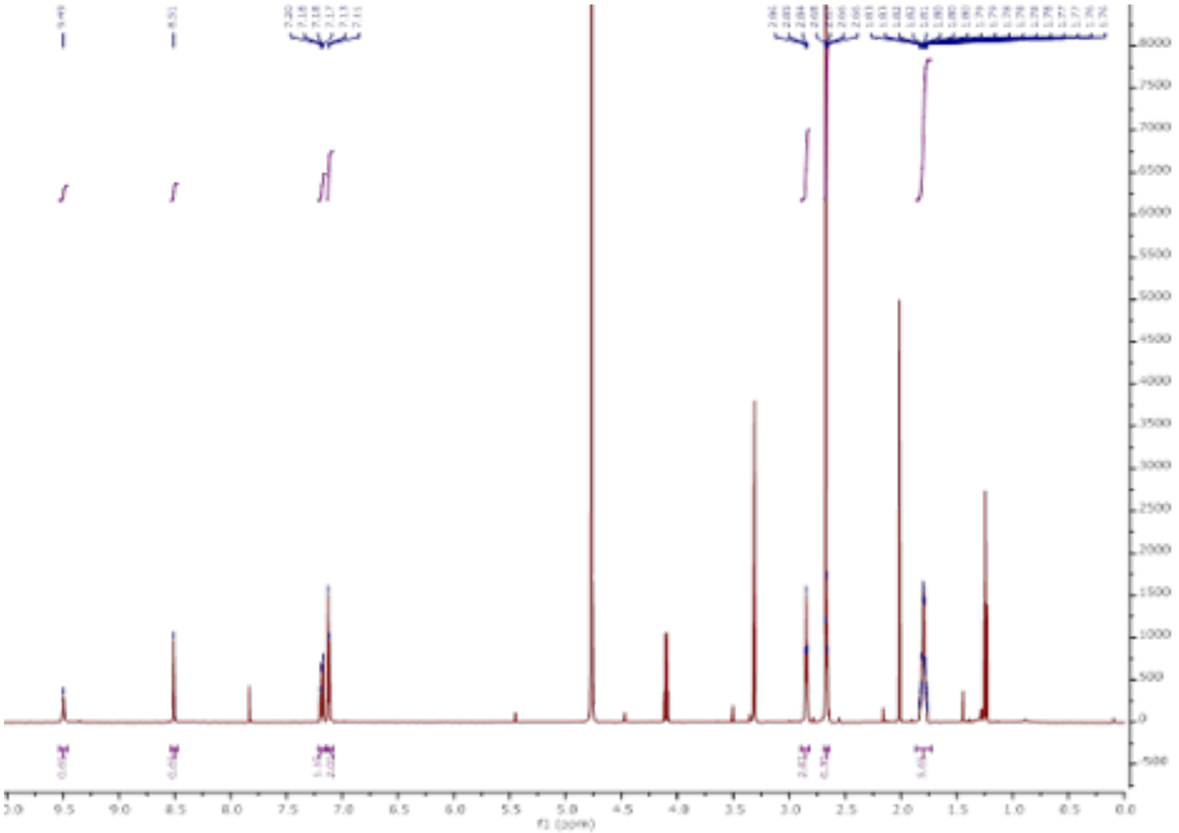

[1]H NMR spectrum of compound **5**

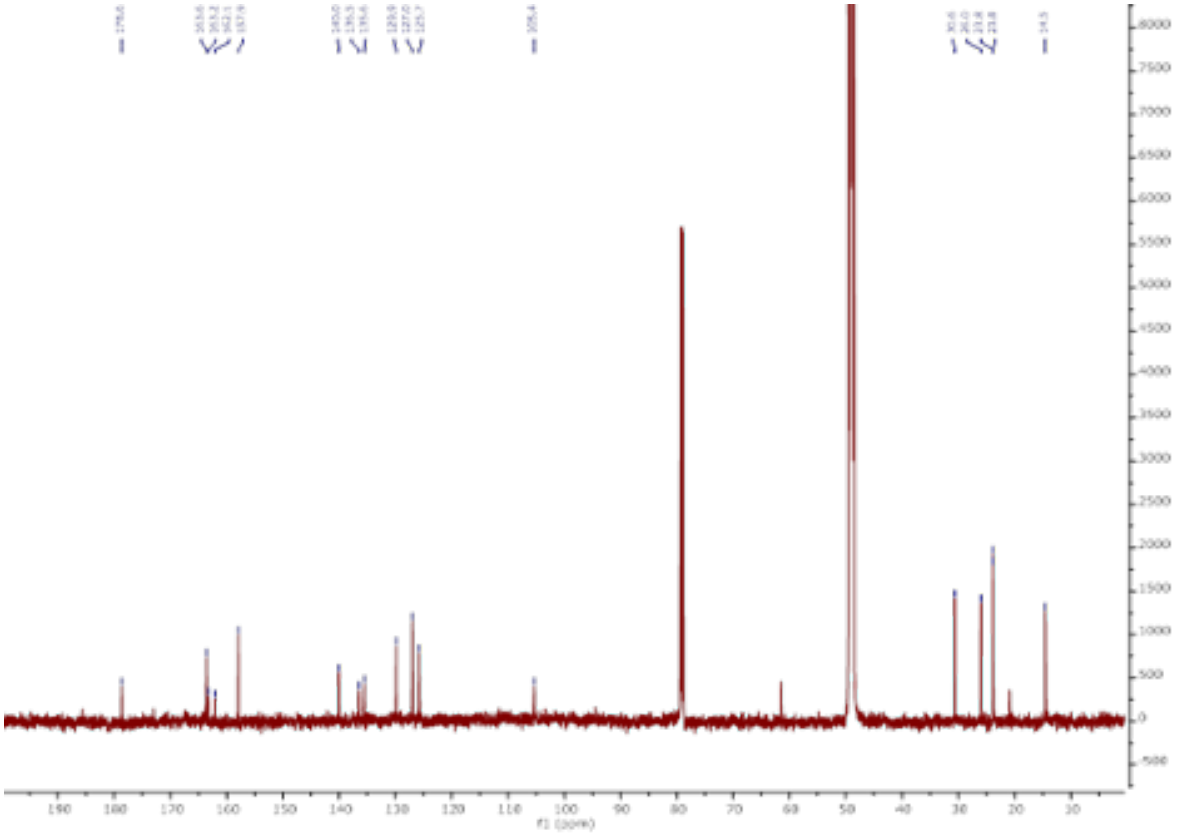

[13]C NMR spectrum of compound **5**

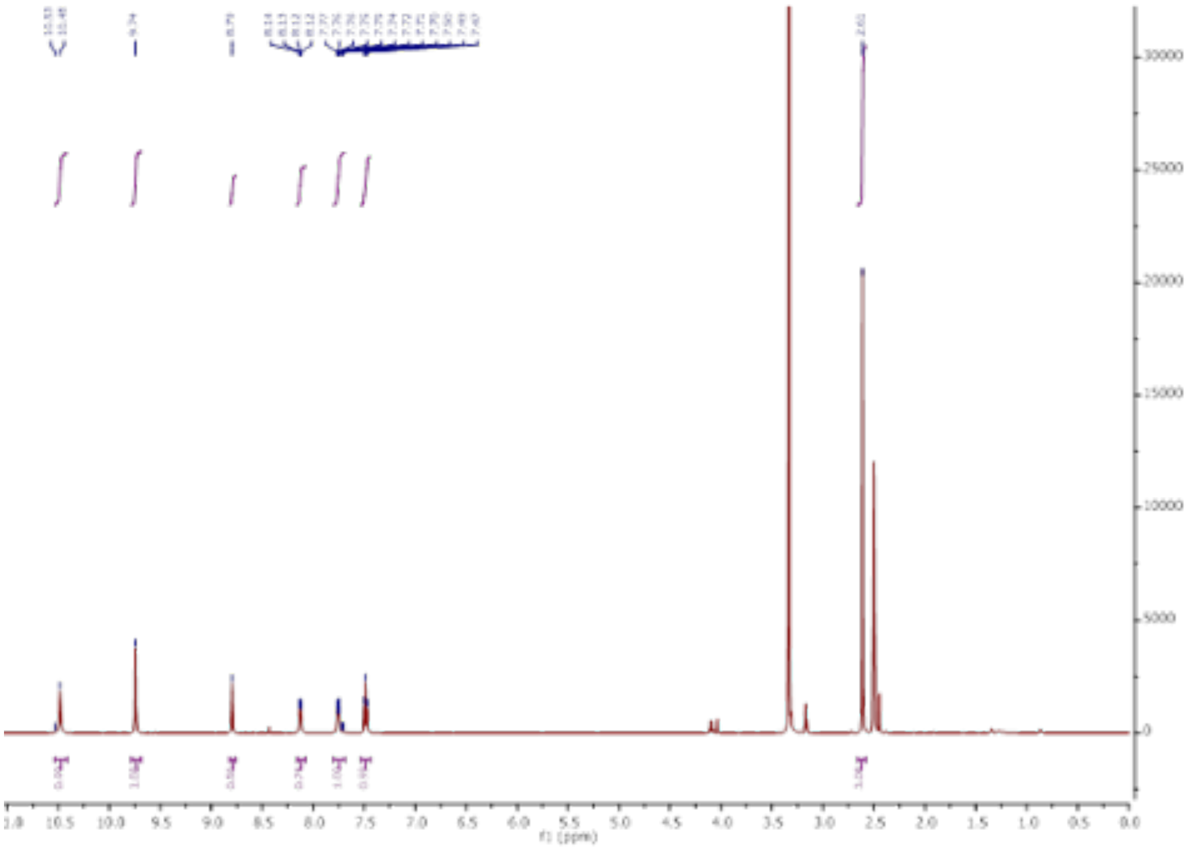

$^{1}H$ NMR spectrum of compound **6**

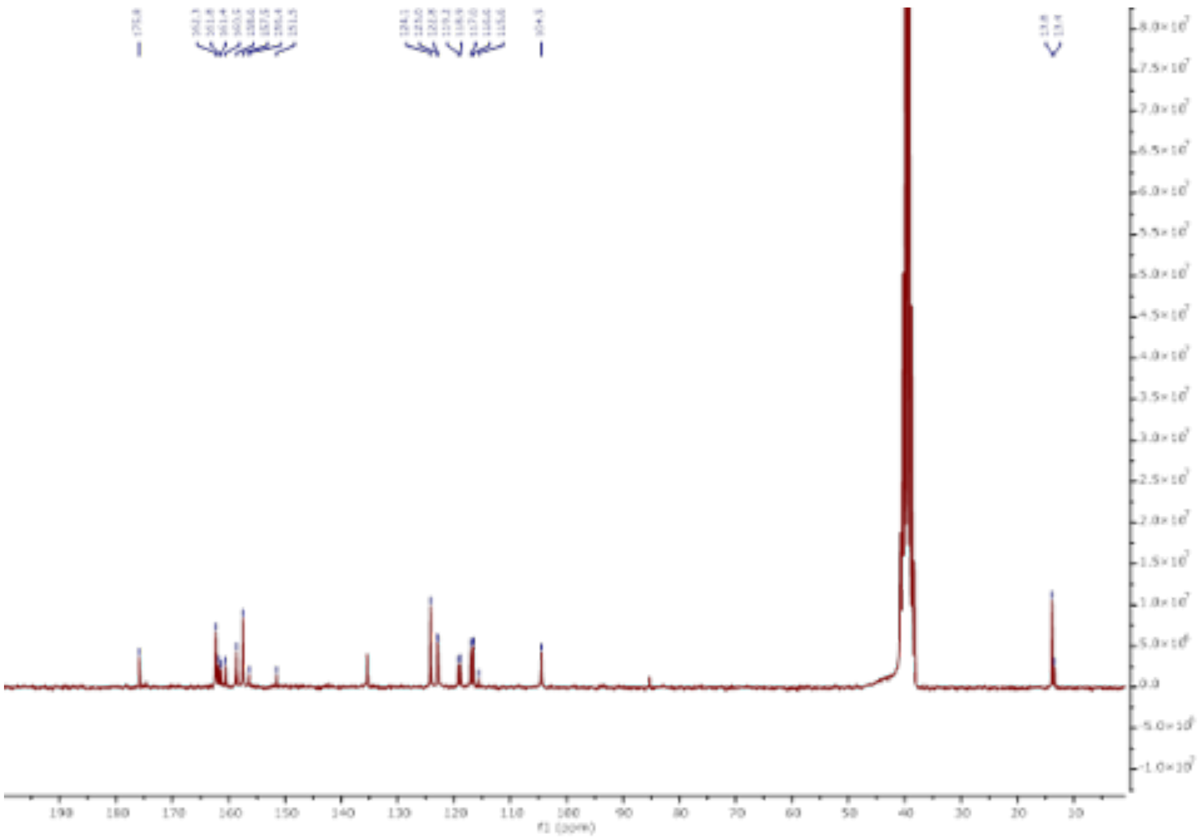

$^{13}C$ NMR spectrum of compound **6**

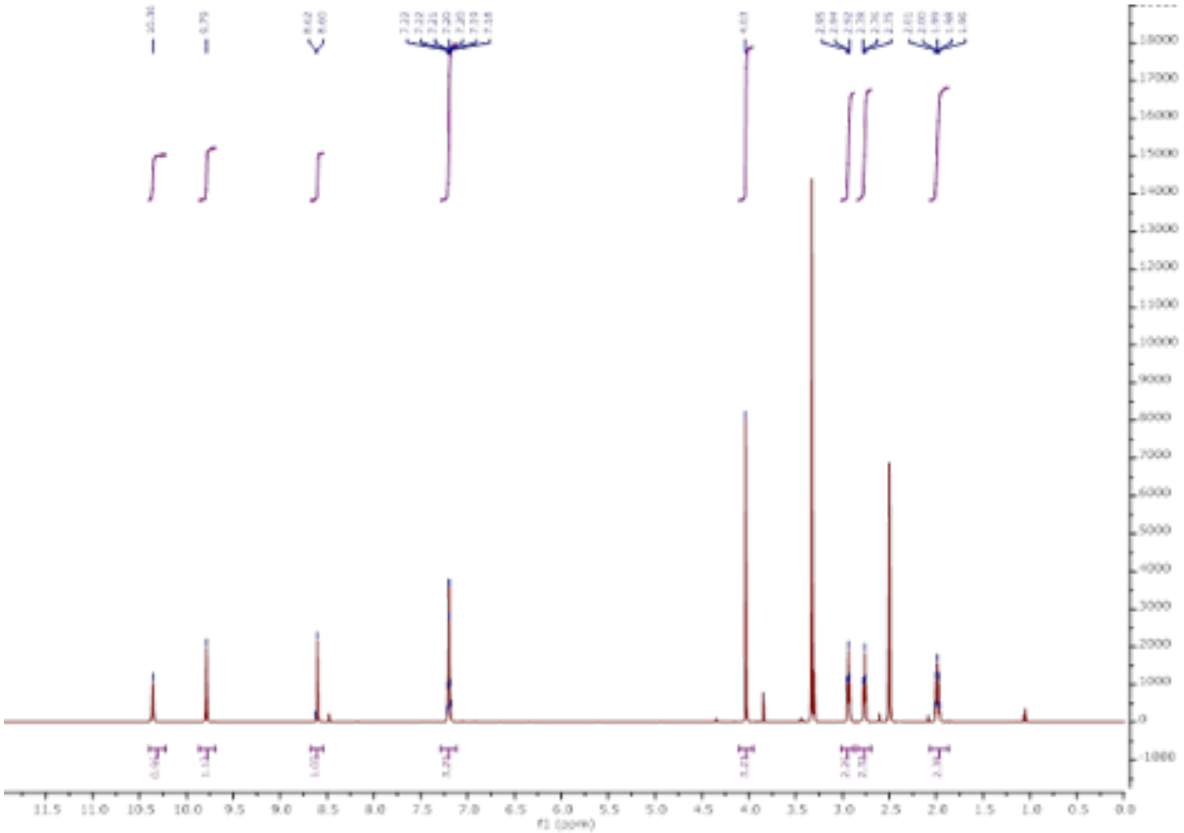

[1]H NMR spectrum of compound **7**

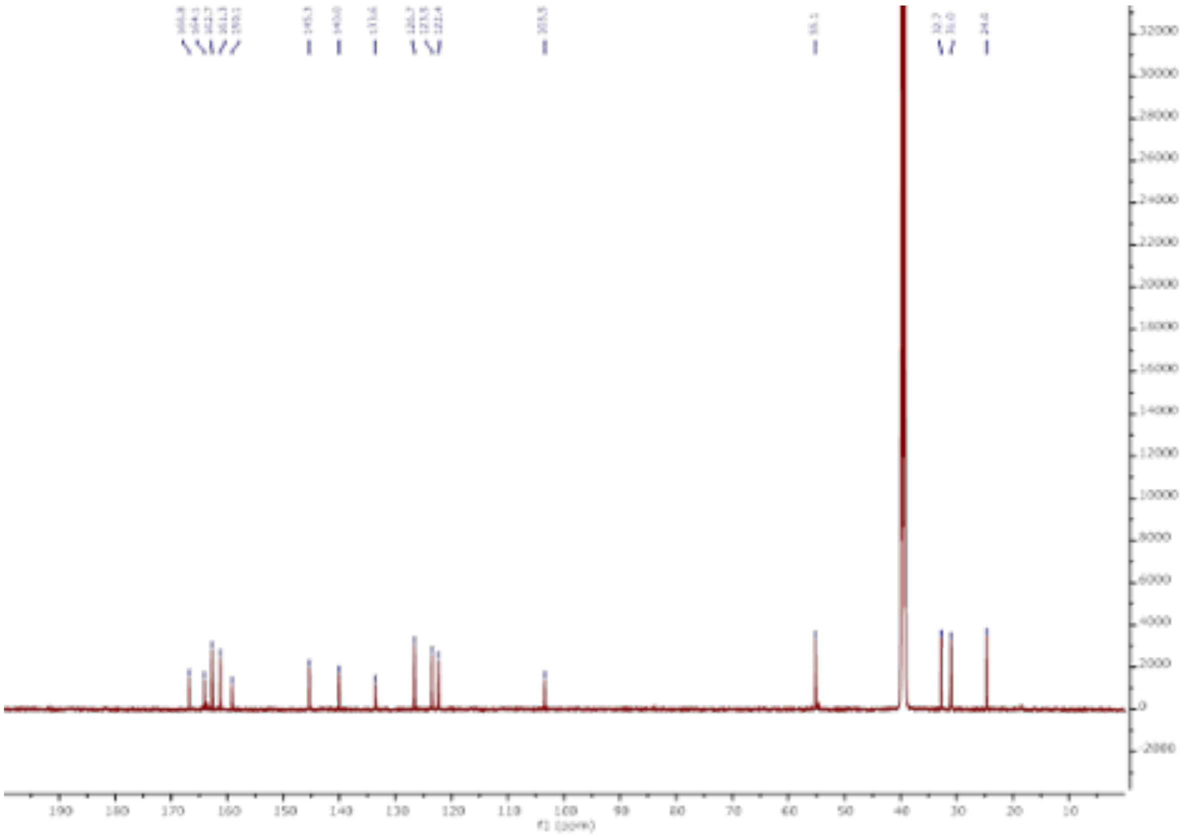

[13]C NMR spectrum of compound **7**

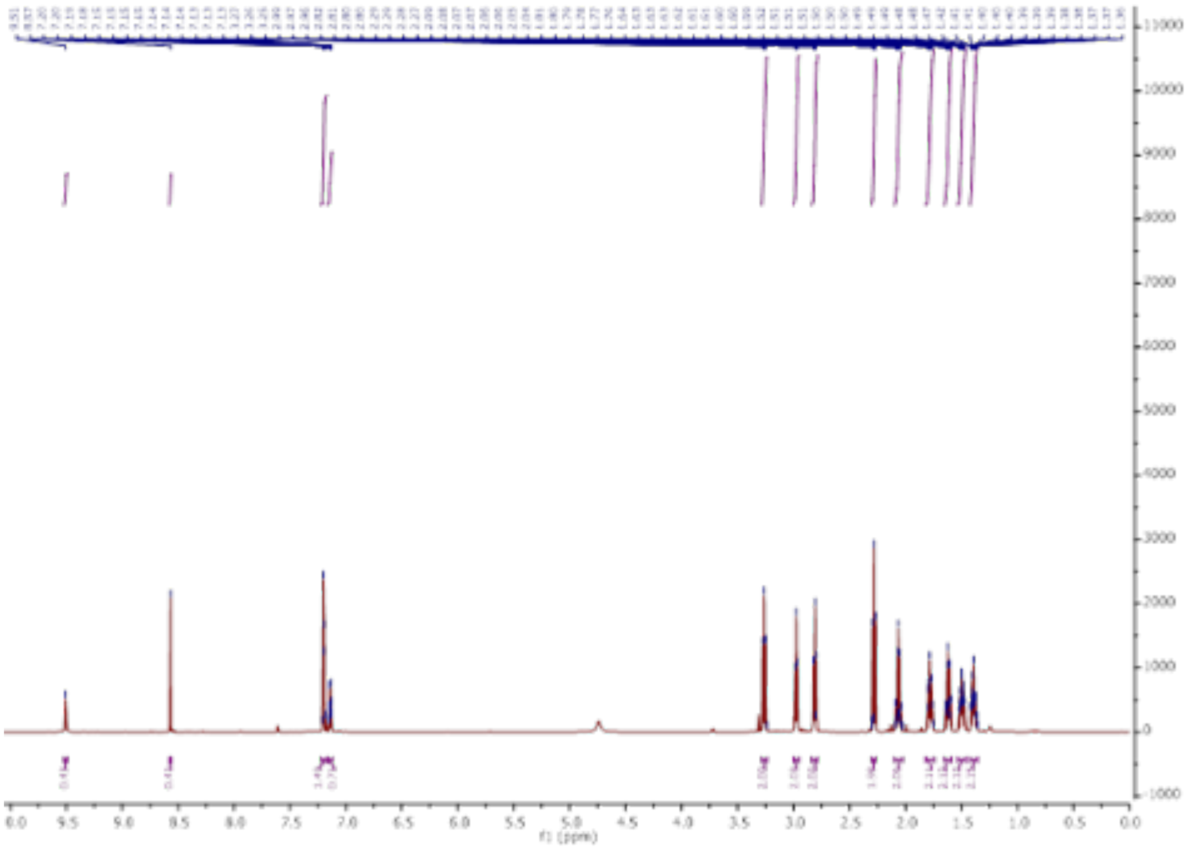

$^{1}H$ NMR spectrum of compound **11**

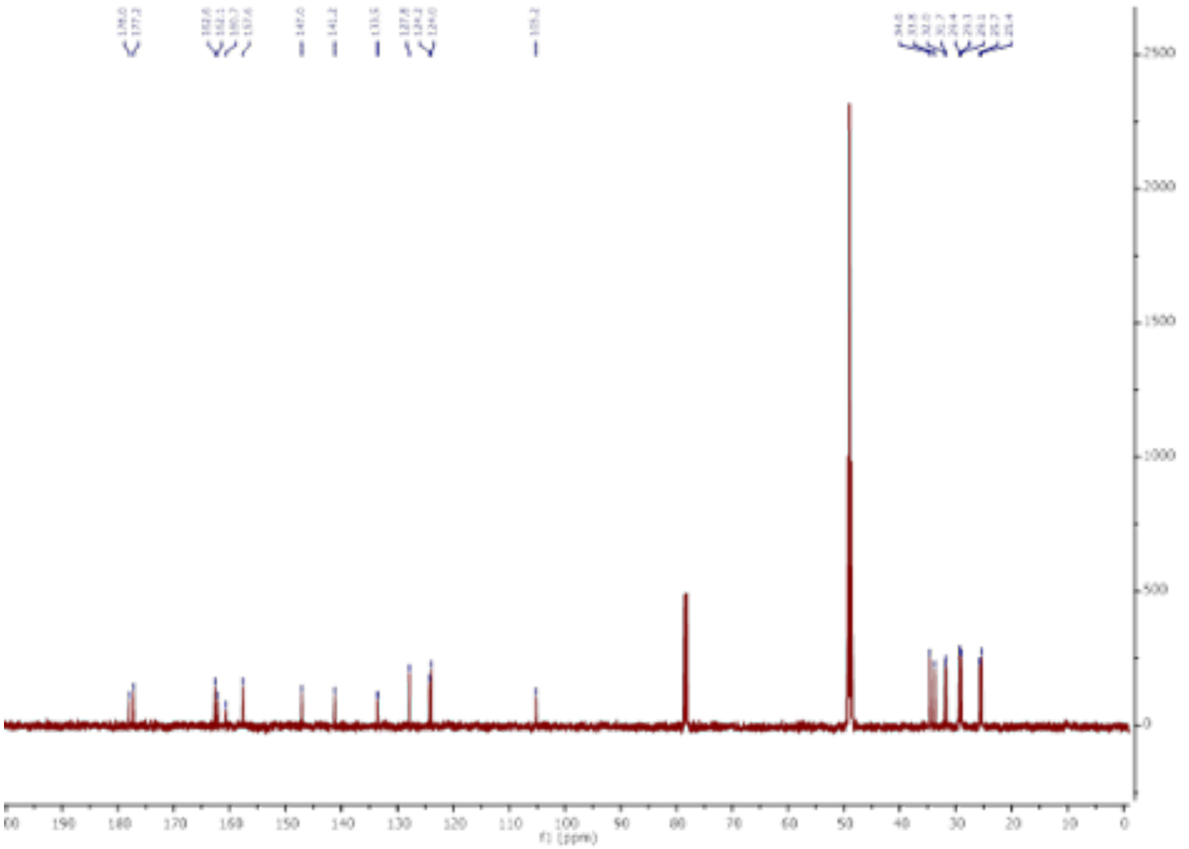

$^{13}C$NMR spectrum of compound **11**

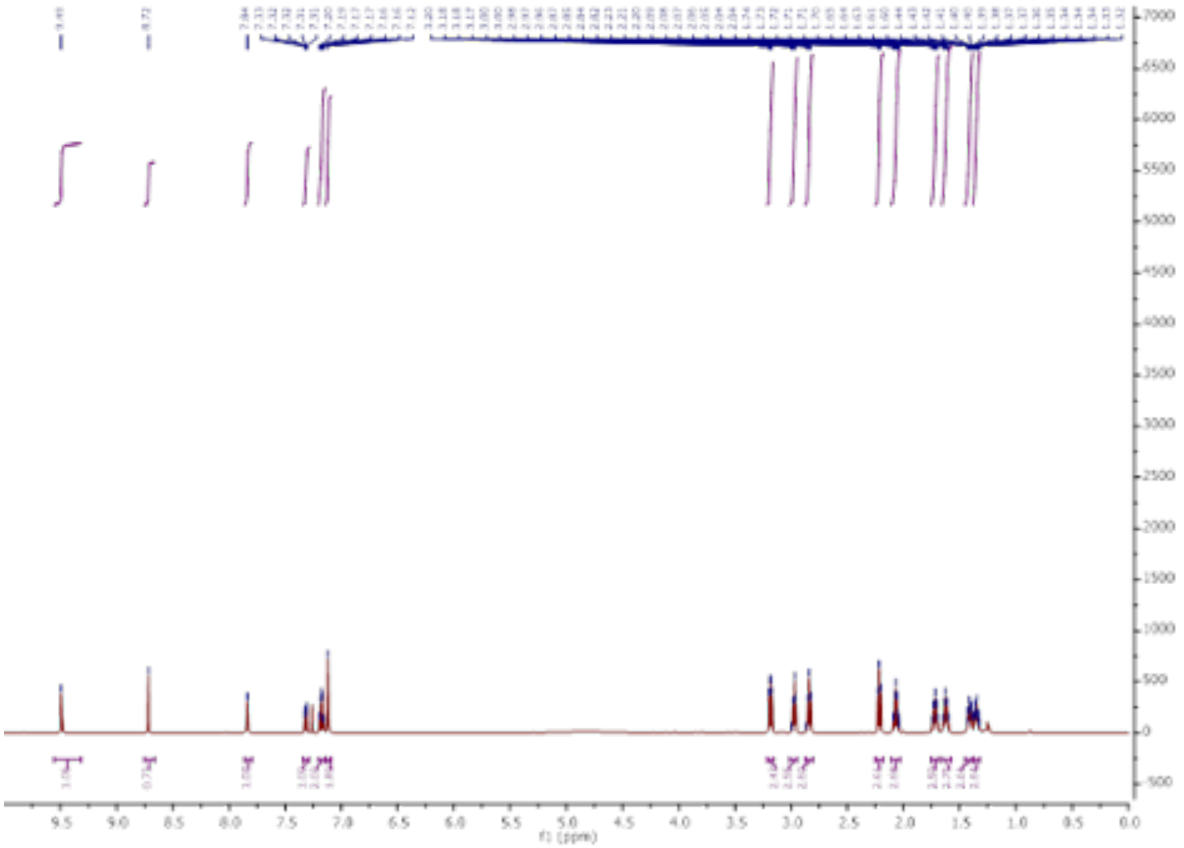

[1]H NMR spectrum of compound **12**

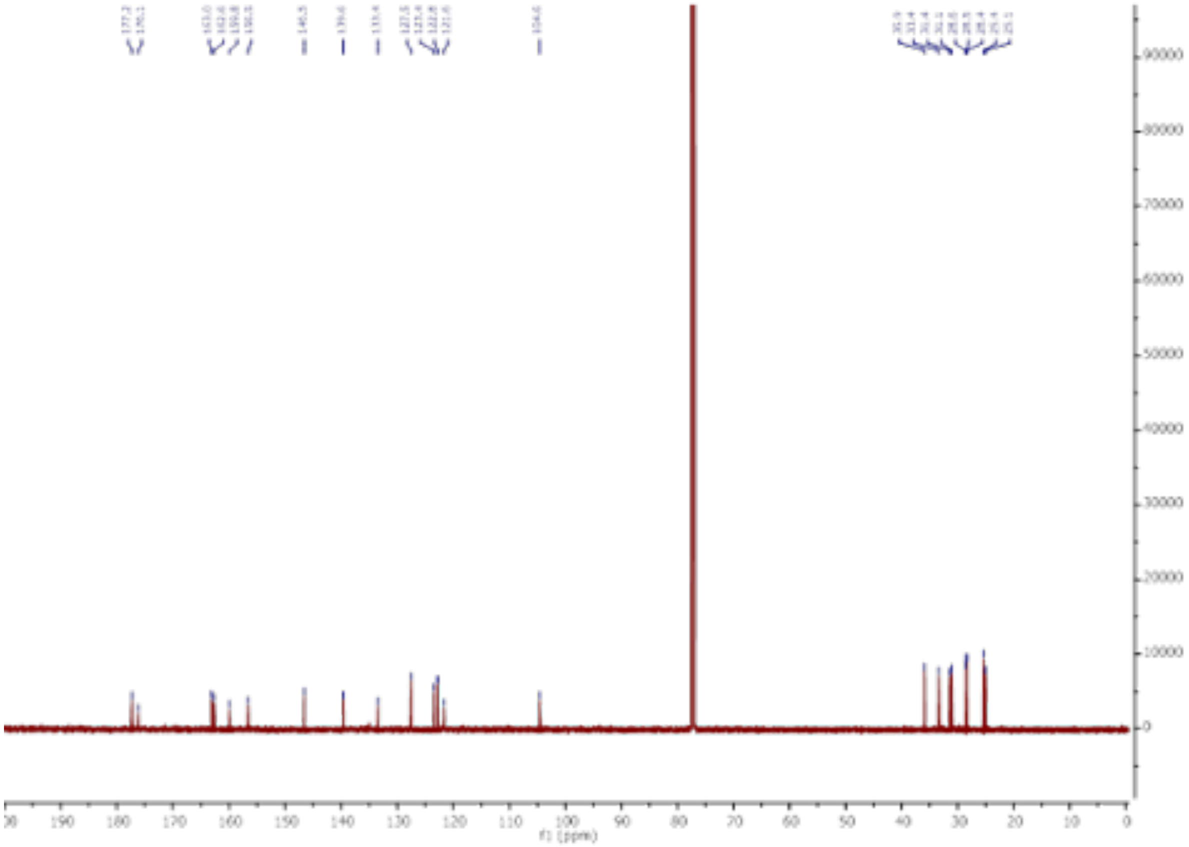

[13]CNMR spectrum of compound **12**

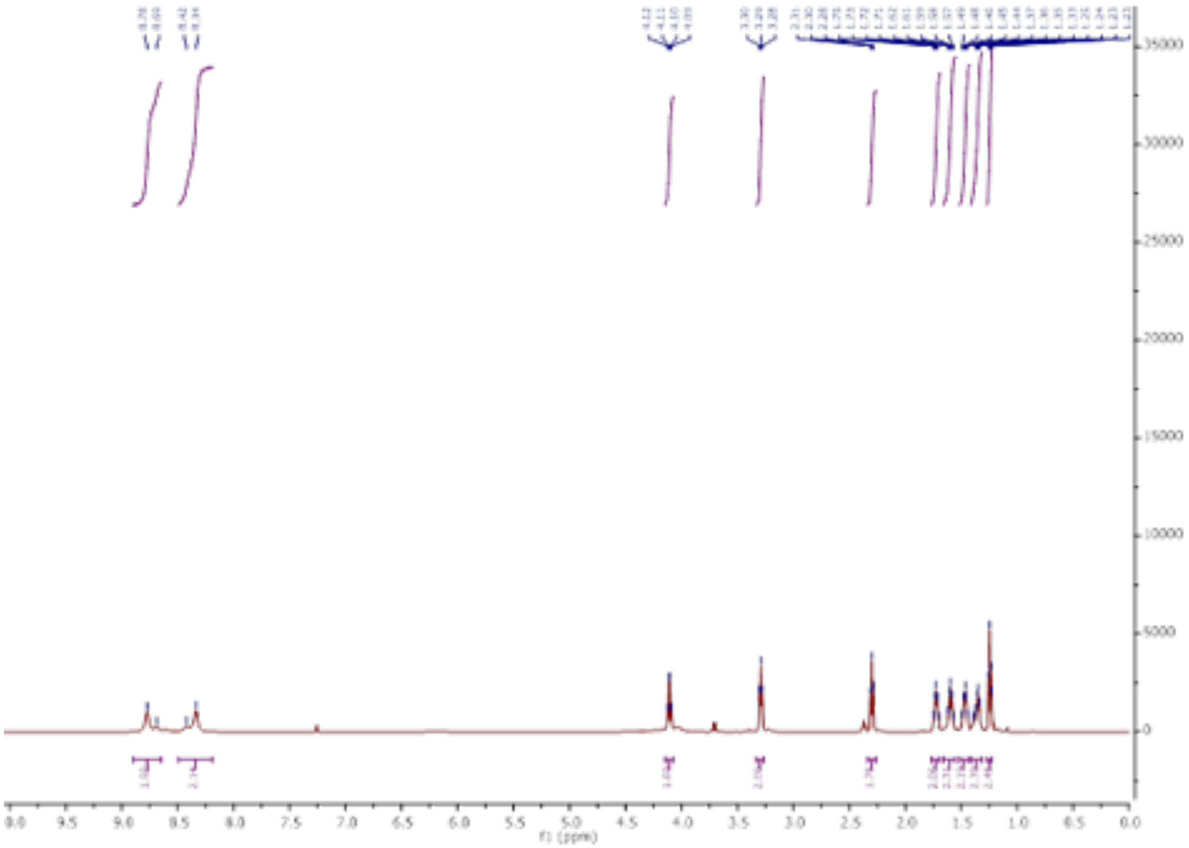

$^{1}$H NMR spectrum of compound **s6**

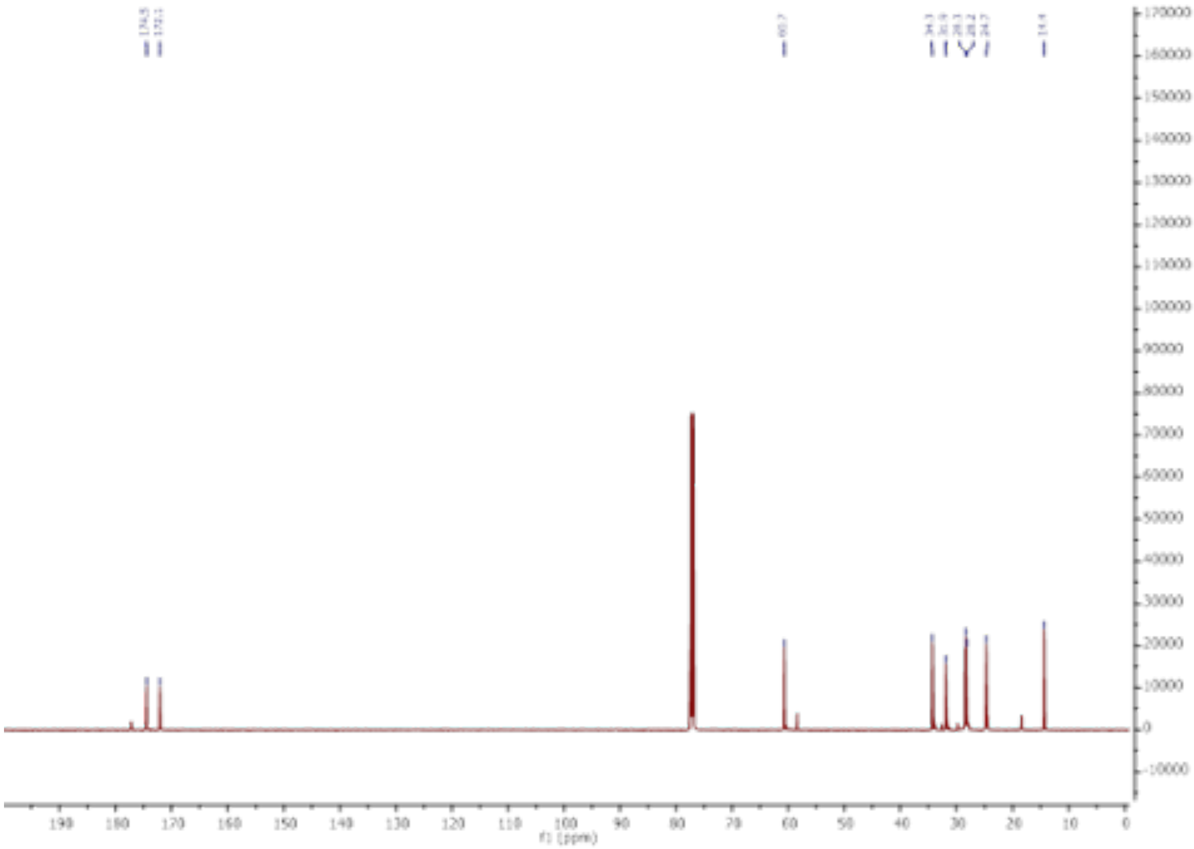

$^{13}$C NMR spectrum of compound **s6**

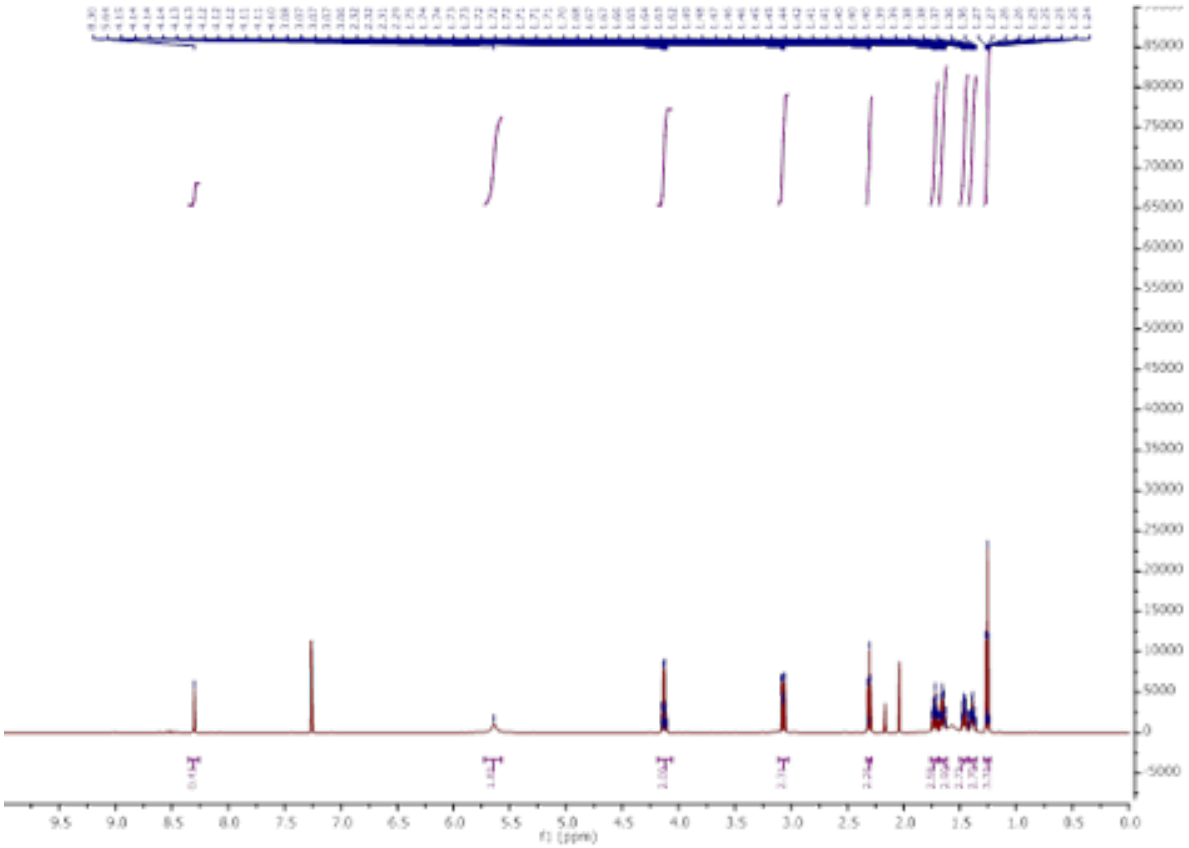

[1]H NMR spectrum of compound **s7**

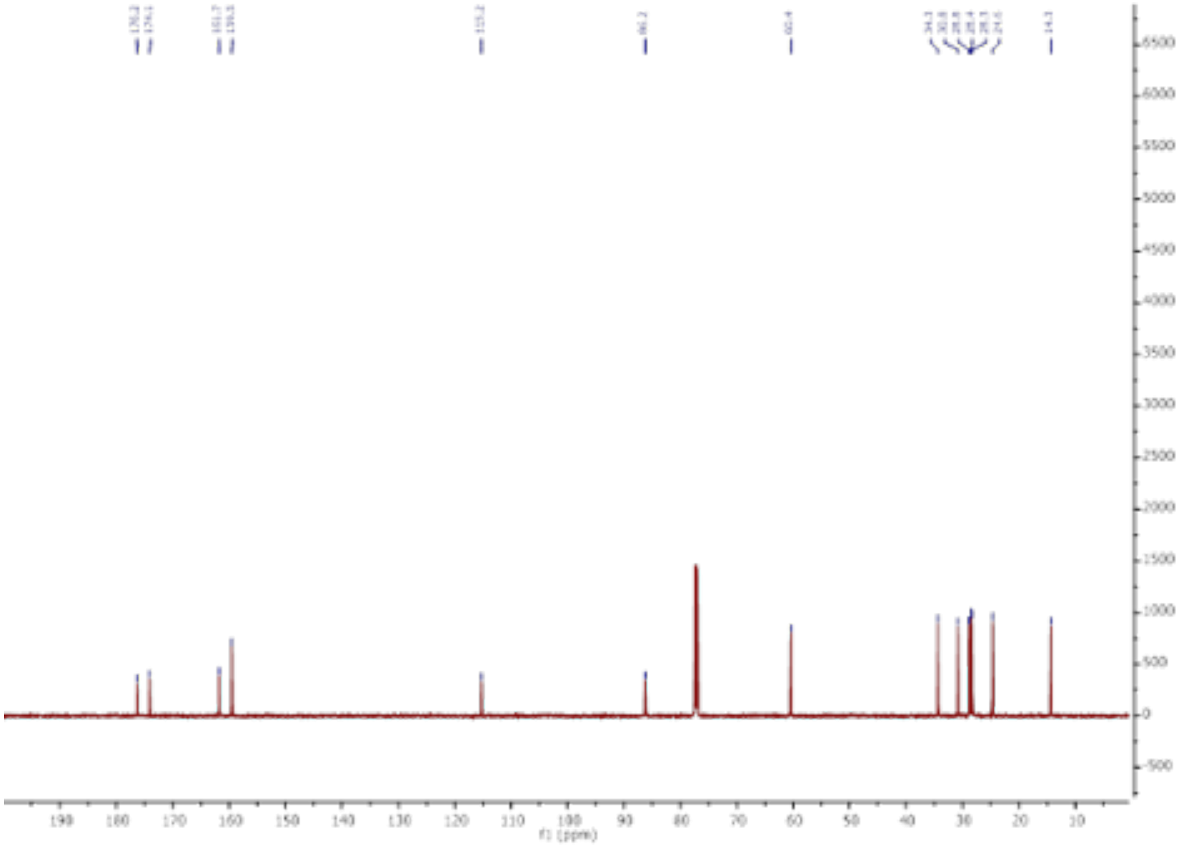

[13]C NMR spectrum of compound **s7**

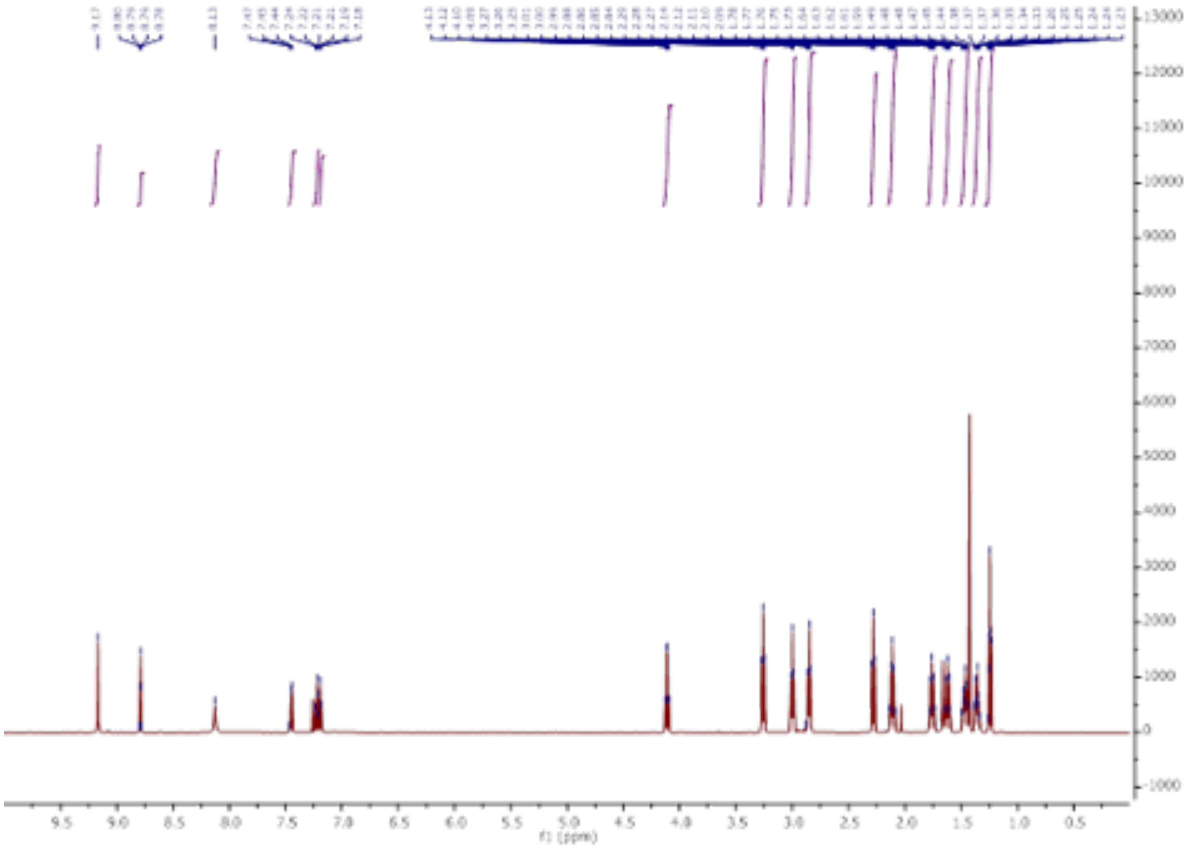

[1]H NMR spectrum of compound **s9**

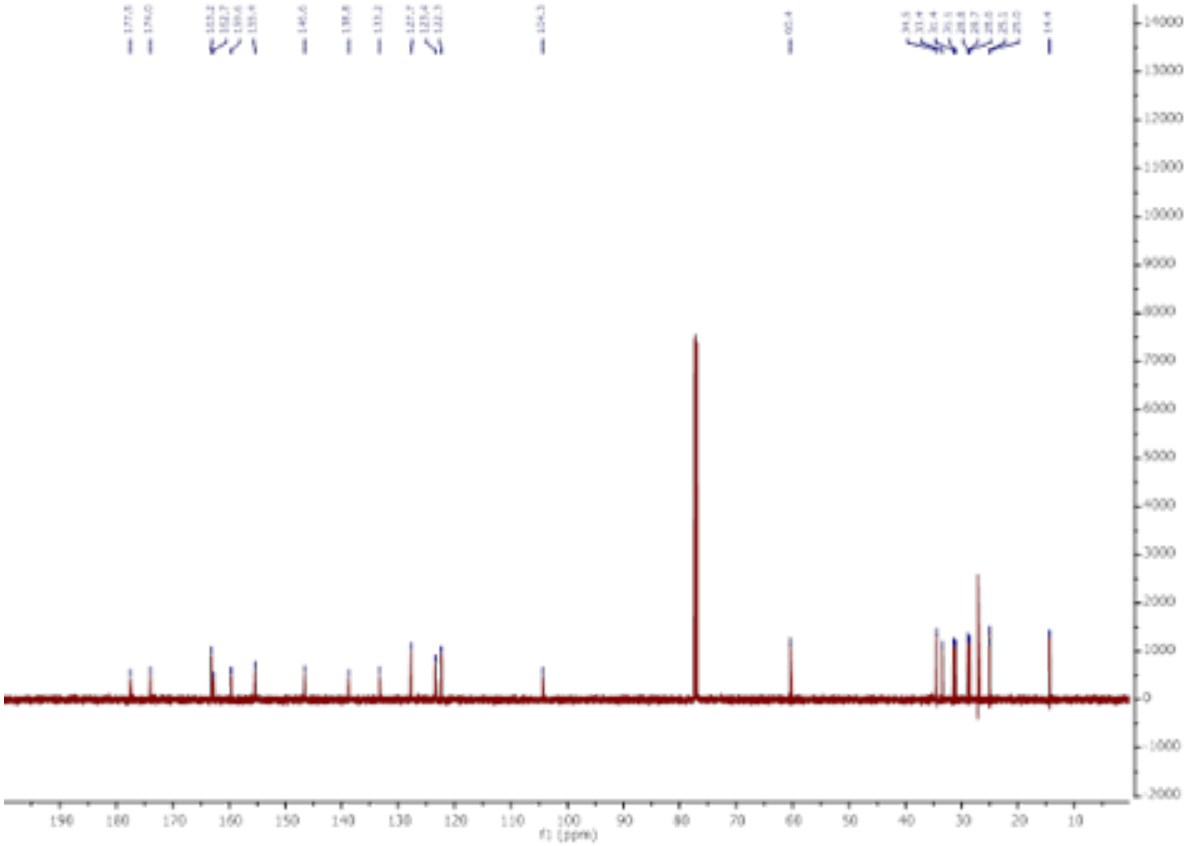

[13]C NMR spectrum of compound **s9**

#### Uncropped gels

##### Figure 3C

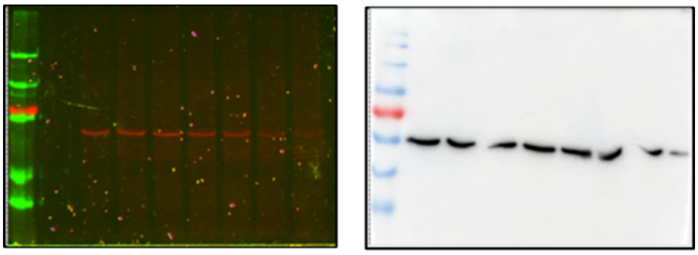

**CEM198 (4) -** Fluorescence

**CEM198 (4) -** anti-Y-TUBA

##### Figure 3D

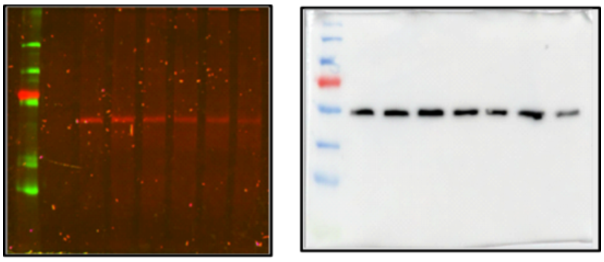

**11 -** Fluorescence

**11 -** anti-Y-TUBA

##### Figure 4A

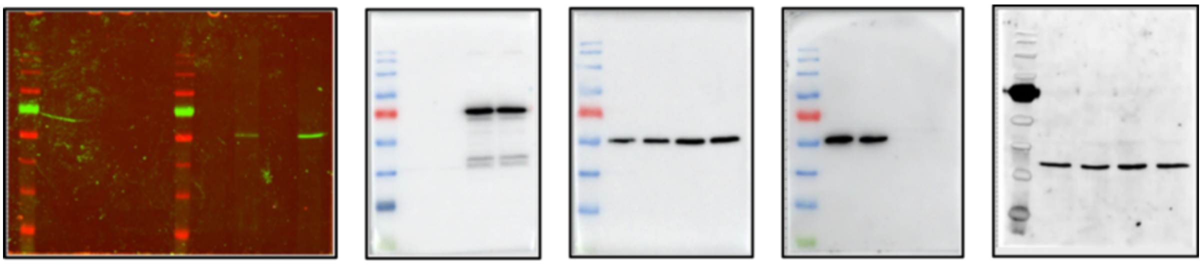

TTL OX -Fluorescence

anti-TTL

anti-Y-TUBA

anti-dY-TUBA

anti-GAPDH

**Figure 4B**

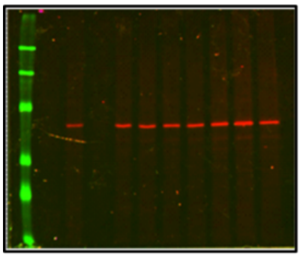

**CEM198 (4) -** Fluorescence with TTL OX

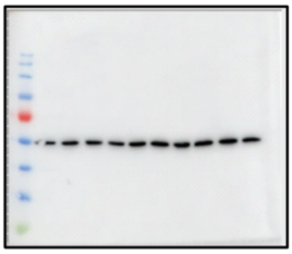

anti-Y-TUBA

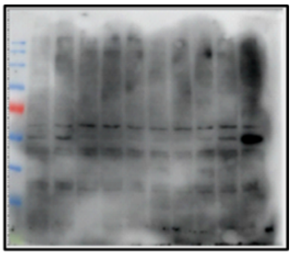

anti-dY-TUBA

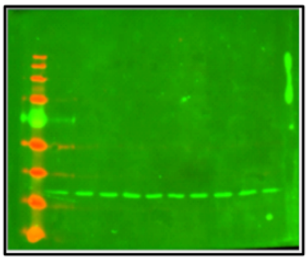

anti-GAPDH

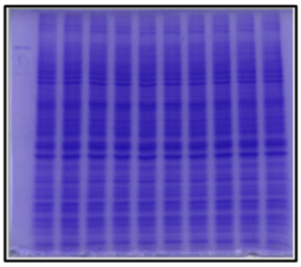

Coomassie stain

**Figure 4C**

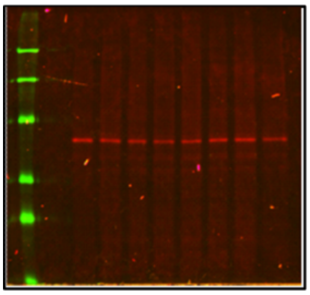

**CEM198 (4) -** Fluorescence without TTL OX

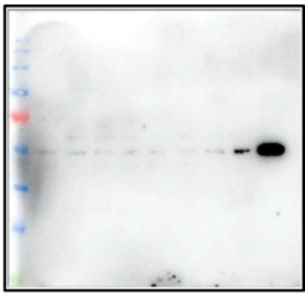

anti-dY-TUBA

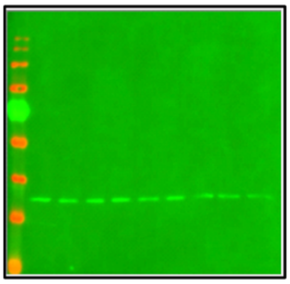

anti-GAPDH

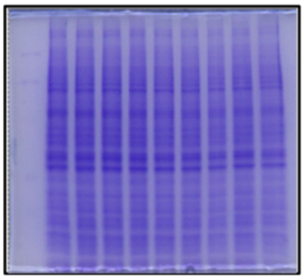

Coomassie stain

**Figure 4D**

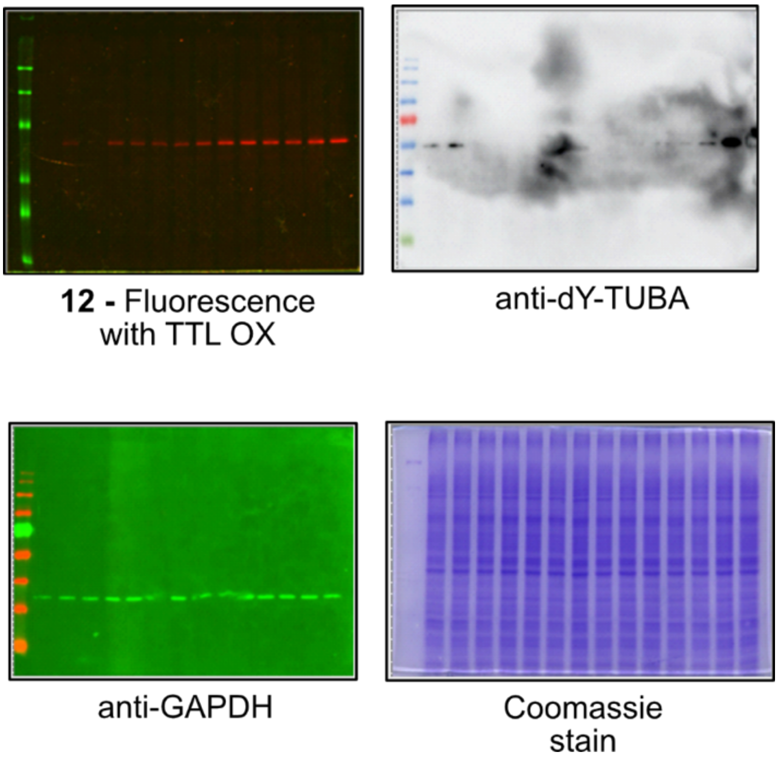

**Figure 4E**

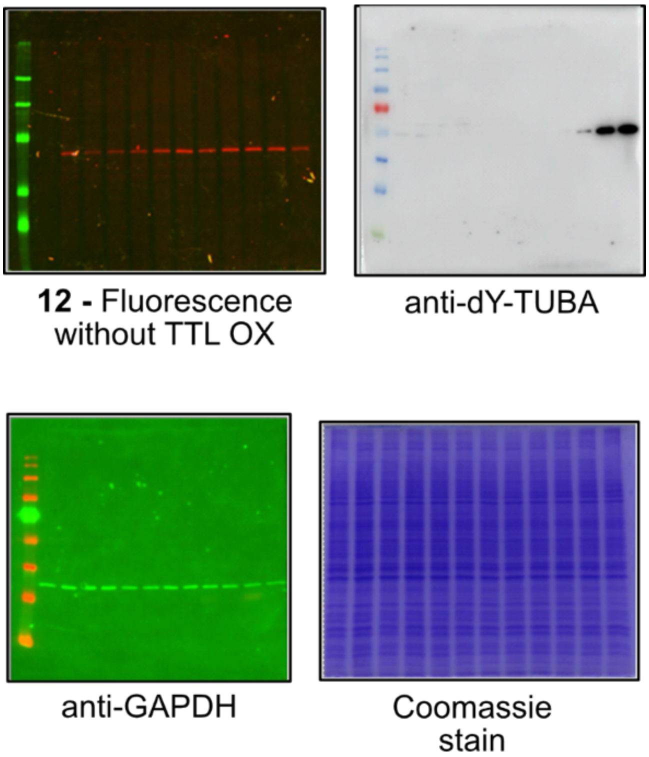

**Figure 5B**

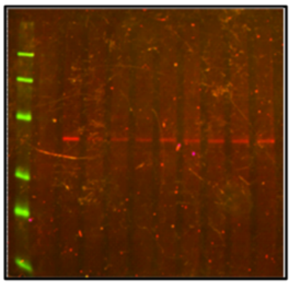

Vinblastin activity

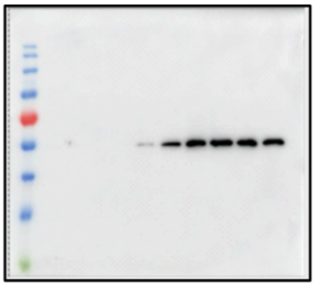

anti-dY-TUBA

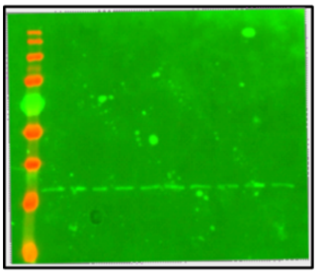

anti-GAPDH

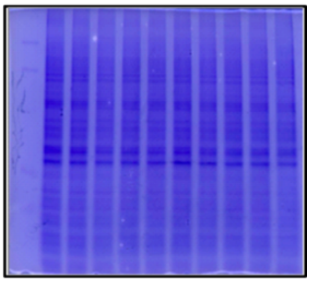

Coomassie
stain

**Figure S1**

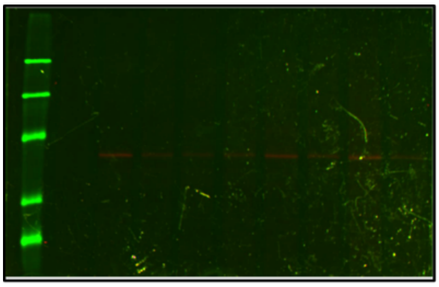

**5** - Fluorescence

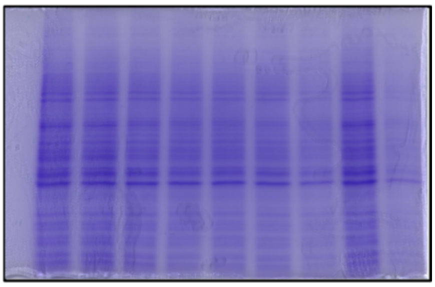

Coomassie
stain

**Figure S2**

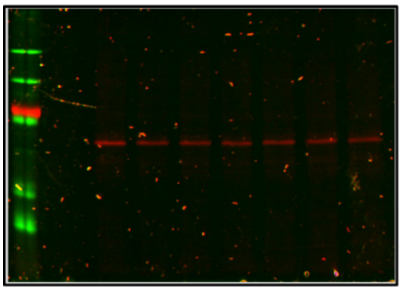

**6** - Fluorescence

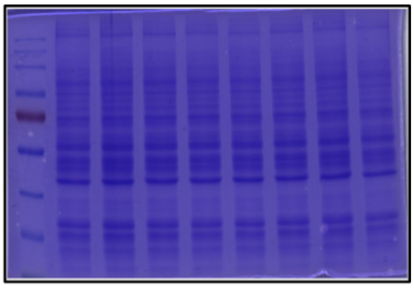

Coomassie stain

**Figure S3**

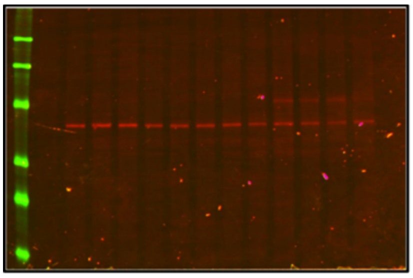

**7** - Fluorescence

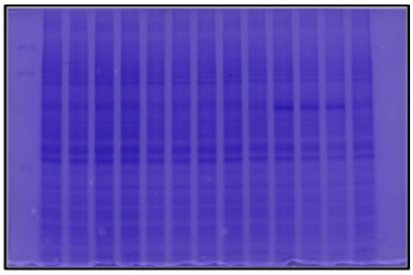

Coomassie stain

**Figure S4**

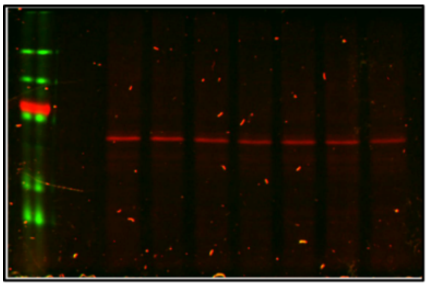

**8** - Fluorescence

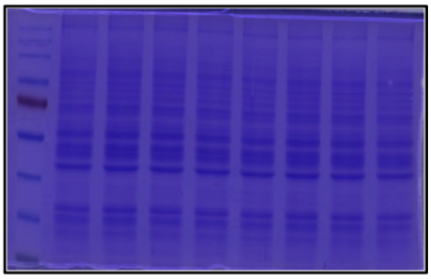

Coomassie stain

**Figure S5**

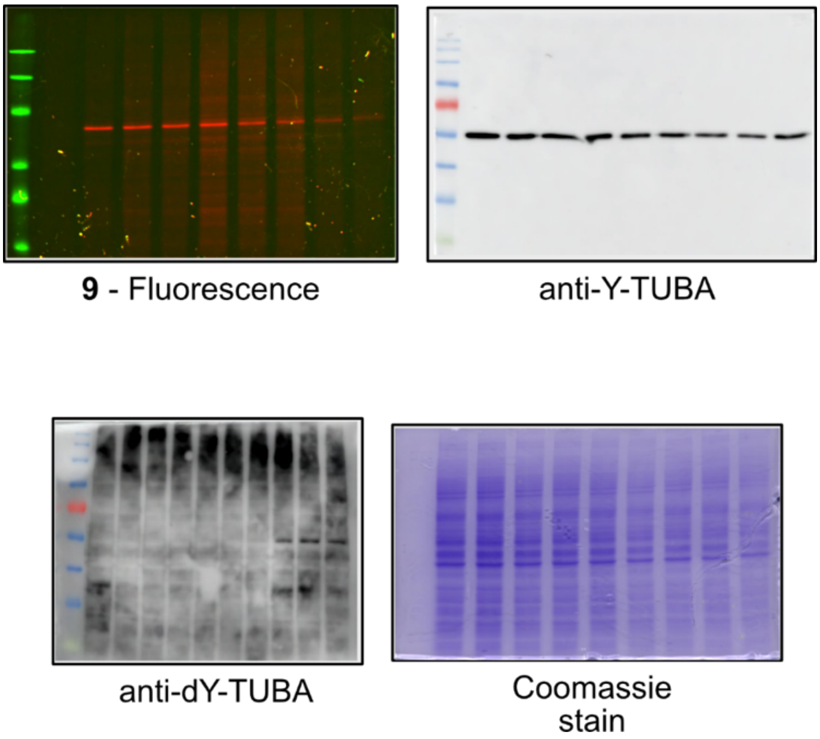

**Figure S6**

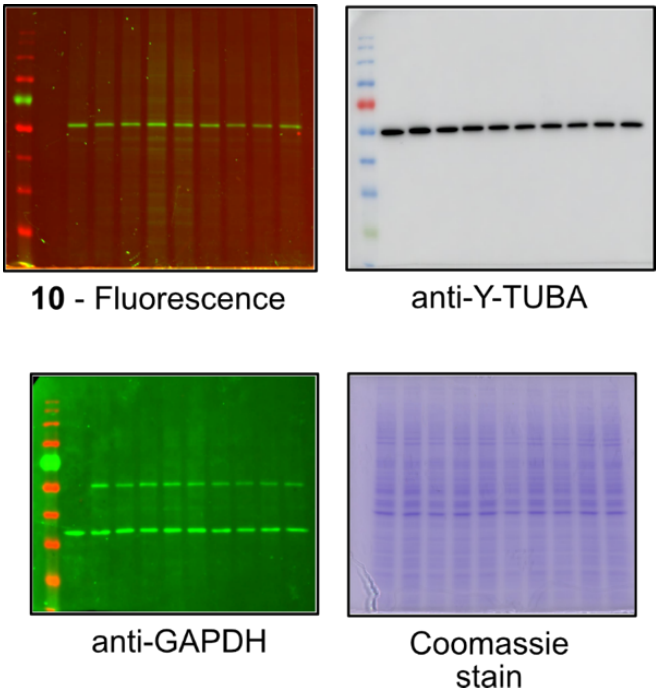

**Figure S8**

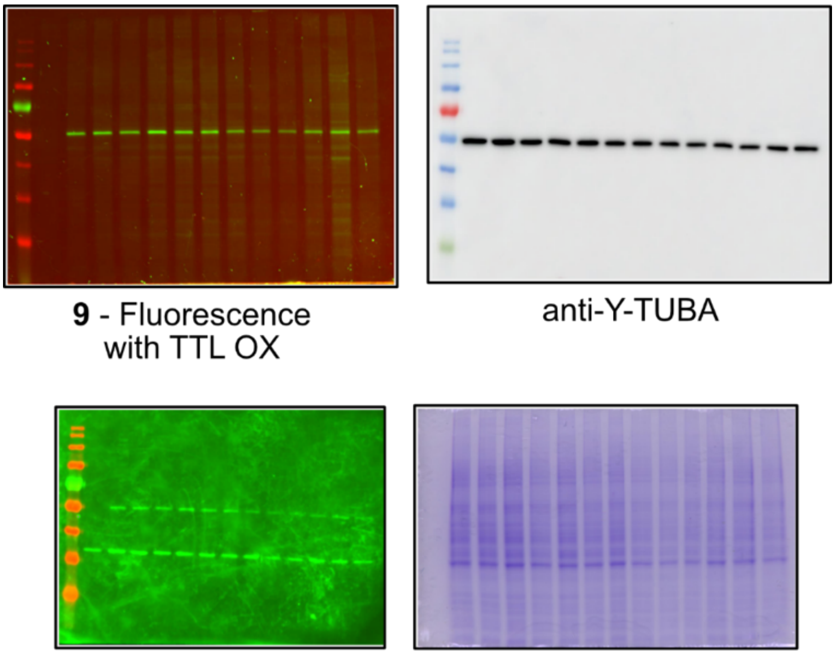

**9** - Fluorescence with TTL OX

anti-Y-TUBA

anti-GAPDH

Coomassie stain

**Figure S9**

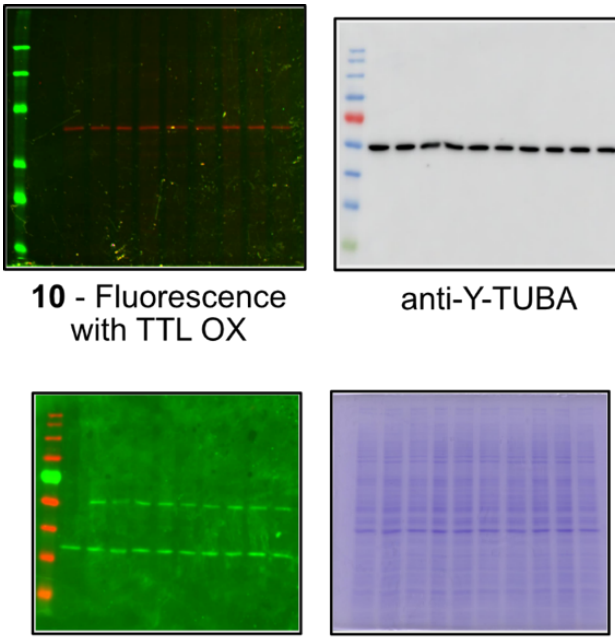

**10** - Fluorescence with TTL OX

anti-Y-TUBA

anti-GAPDH

Coomassie stain

**Figure S10**

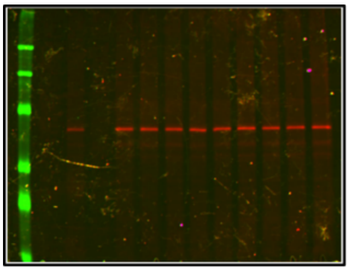

**11** - Fluorescence with TTL OX

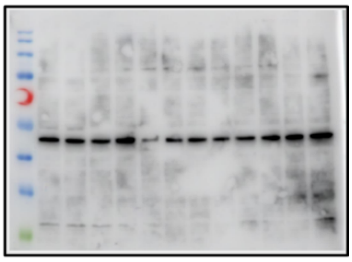

anti-dY-TUBA

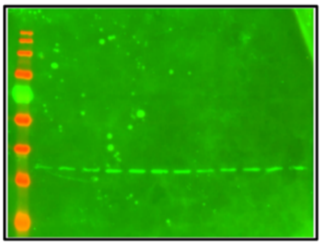

anti-GAPDH

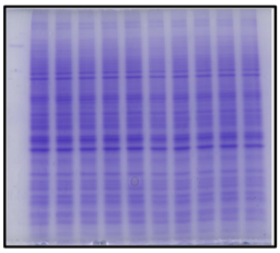

Coomassie stain

**Figure S11**

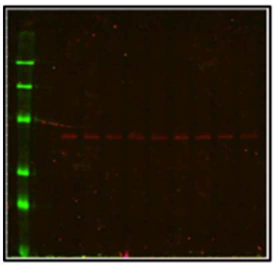

Colchicine activity

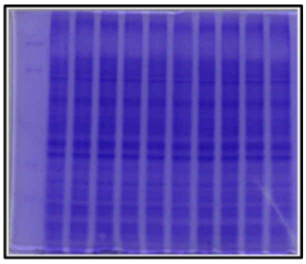

Coomassie stain
